# Large population sizes shape genome evolution in barnacles

**DOI:** 10.64898/2026.04.27.721231

**Authors:** Angel G. Rivera-Colón, Scott T. Small, Erin Jezuit, John P. Wares, Andrew D. Kern

## Abstract

Population size is a key factor underlying the mode and tempo of evolution, particularly as it relates to the strength of selection and drift. While the mechanisms underlying the interactions between population size and selection have been studied in population genetics for over a century, empirical knowledge of these dynamics has been limited to species with small-to-moderate historical population sizes. This gap in knowledge highlights the need for empirical studies in systems with historically large population sizes and elevated diversity. The Pacific acorn barnacle (*Balanus glandula*) presents an exceptional model to study evolution in extremely large populations, exhibiting census sizes often exceeding the tens of thousands of individuals per square meter. We generated a new chromosome-level genome assembly for this species, which we leverage in one of the first large-scale genomic analyses in this system. This assembly reveals a highly polymorphic genome with over 3% heterozygosity. At a population level, *B. glandula* exhibits extreme levels of polymorphism, with nucleotide diversity surpassing 5% genome-wide in just a small collection of individuals. Across the genome, nucleotide diversity predictably decreases at functional elements, including both coding and non-coding sequences, likely reflecting strong purifying selection along the genome. At the same time, McDonald-Kreitman tests reveal that the majority of non-synonymous substitutions between barnacle species were driven by positive selection, consistent with the expected increase in the efficacy of selection in large populations. These remarkable levels of diversity set *B. glandula* as a unique model for the study of evolution at extreme demographic scales and highlights the importance of testing evolutionary theory across a wide variety of empirical systems.

## Introduction

The size of a population is a key factor mediating the mode and tempo of evolution, particularly as it relates to the strength of stochastic versus deterministic evolutionary processes (J. H. Gillespie, 2000; Kimura et al., 1963). Population genetics theory predicts that selection is less efficacious in small populations (Charlesworth, 2009; Gravel, 2016), resulting in reduced genetic diversity from genetic drift and an increase in genetic load due to the accumulation of deleterious mutations (Dussex et al., 2023; Lande, 1994; Lynch et al., 1995). These dynamics have been extensively observed in natural populations, e.g., the increase in genetic load in humans as a function of their geographic distance to more diverse groups in sub-Saharan Africa (Henn et al., 2016) and the increase in genetic load and inbreeding depression in small populations of conservation concern (Frankham, 2005; Kardos et al., 2023; Lande, 1988; Westemeier et al., 1998), empirically supporting the theoretical expectations.

In contrast, evolution in large populations is thought to be primarily mediated by deterministic processes due to an increase in the efficacy of selection (Charlesworth, 2009; Gravel, 2016; Lanfear et al., 2014). The dynamics of selection in large populations are expected to produce patterns such as clonal-interference effects driven by soft sweeps and recurrent mutation (Park & Krug, 2007; Weissman & Hallatschek, 2014), as well as the loss of genetic diversity through the hitchhiking effect (J. M. Smith & Haigh, 1974). In sufficiently large populations, this process is predicted to be a primary driver of stochastic diversity loss (J. H. Gillespie, 1999, 2000). Yet, while there have been empirical studies characterizing genetic diversity and the mechanisms of evolution in large populations (e.g., Anderson et al., 2017; Brasó-Vives et al., 2026; Dey et al., 2013; Teterina et al., 2025; The *Anopheles gambiae* 1000 Genomes Consortium, 2017), they remain comparatively rare. Empirical work on this subject has mainly focused on eukaryotic species with small-to-moderate historical population sizes (Buffalo, 2021; Cutter et al., 2013; Leffler et al., 2012), limiting our capacity to observe these evolutionary dynamics at demographic extremes. This gap highlights the need for empirical studies in systems with historically large population sizes and elevated levels of genetic polymorphism.

The Pacific acorn barnacle (*Balanus glandula*; Balanidae) presents an exceptional model for studying the dynamics and mechanisms of evolution in large populations. This species is widespread across the Pacific intertidal of North America, from Baja California to Alaska, where it can reach densities of 10^4^ of individuals per square meter (Menge, 2000). Like most acorn barnacles, *B. glandula* is a sessile, simultaneous hermaphrodite that reproduces through cross-fertilization, mating with its neighbors via pseudo-copulation; even isolated individuals, beyond the reach of any mate, can produce outcrossed broods by capturing water-borne sperm (Barazandeh et al., 2014). Individuals of this species can be long-lived (8–10 years; Newman and Abbott, 1980) and show high fecundity, with 2–6 broods per year (Hines, 1978; Høeg et al., 1987), brood sizes of 10^3^–10^4^ embryos (Barnes & Barnes, 1956), and a reproductive output of *≈* 10^3^ nauplius larvae per individual (Leslie et al., 2005). Additionally, *B. glandula* displays high dispersive potential. Nauplius larvae feed and develop in the plankton for a period of weeks before settling (Brown & Roughgarden, 1985; Pfeiffer-Hoyt & McManus, 2005), leading to their dispersal up to hundreds of kilometers (Sotka & Palumbi, 2006; Sotka et al., 2004). This combination of high fecundity, outcrossing, and long-lived planktonic larvae capable of long-distance dispersal, together with these enormous census sizes, is consistent with very large effective population sizes, making *B. glandula* an ideal system for testing theoretical predictions about selection and drift in large populations.

This species has been the focus of extensive ecological and genetic research. Prior work has linked environmental factors to variation in development, recruitment, dispersal, and reproduction (Barshis et al., 2011; Berger, 2009; Berger et al., 2006; Pfeiffer-Hoyt & McManus, 2005; Schubart et al., 1995), and substantial effort has characterized the spatial population structure of *B. glandula* along the Pacific coast, particularly a steep genetic cline on the coast of California that separates northern and southern populations (Sotka et al., 2004; Wares & Cunningham, 2005; Wares & Skoczen, 2019; Wares et al., 2021). Yet, *B. glandula* remains genomically understudied. Previous research has been limited to a handful of nuclear and mitochondrial markers, with the most extensive study employing reduced representation sequencing in the absence of a reference genome (Wares et al., 2021). Moreover, while two reference assemblies for *B. glandula* exist in public databases, both are highly fragmented and likely incomplete, which limits their use when studying genome-wide patterns of diversity and selection in this system.

Most publicly available barnacle assemblies suffer from similar limitations, presumably due to the difficulty of assembling highly polymorphic genomes from short reads (Kajitani et al., 2014; Vinson et al., 2005). Despite this limitation, a handful of chromosome-level assemblies have been recently generated for several barnacle species (Bernot et al., 2022; Bishop et al., 2025; Blaxter et al., 2023; B. J. Carlson et al., 2025; Han et al., 2024). Yet, these genomes largely belong to evolutionarily or geographically distant species. While useful for the long-term comparisons we perform here, their evolutionary distance limits the utility of these resources for studies in *B. glandula*. This drives the need for high-quality assemblies of both evolutionarily closely related species, which could serve as outgroups for recent evolutionary comparisons, and co-localized species, which enable the study of ecological factors maintaining diversity in this system.

Work in other barnacle species has revealed substantial levels of genetic diversity (Alm Rosenblad et al., 2021), in accordance with the expectation of very large effective populations, and complex patterns of selection at remarkably small spatial scales. The best-characterized example is the balanced polymorphism at the mannose-6-phosphate isomerase (*Mpi*) locus in the northern acorn barnacle *Semibalanus balanoides*, which segregates across intertidal environments (Nunez et al., 2020; Schmidt et al., 2000). Building on this single-locus work, whole-genome pooled resequencing across intertidal microhabitats has shown that spatially varying selection extends beyond *Mpi*, leaving genomic signatures of adaptation to environmental heterogeneity at loci throughout the *S. balanoides* genome (Nunez et al., 2021). These studies demonstrate that selection can structure barnacle genetic variation over just meters of tidal height; however, such genome-wide inference has so far relied on a fragmented draft assembly in a single species, and the landscape of diversity and selection remains uncharacterized across most of the barnacle tree, including *B. glandula*.

In this work we start addressing the need for empirical systems to study evolution in large populations by generating a high-quality, chromosome-level assembly for the Pacific acorn barnacle. We additionally generate a draft assembly for its congener, the wrinkled barnacle *Balanus crenatus*, providing a closely related outgroup for divergence-based analyses. We use these assemblies to perform whole-genome comparative analyses, revealing conservation in genome architecture across 165 million years of barnacle evolution. We further conduct the first genome-wide assessment of genetic variation in this system, uncovering extreme levels (*>* 5%) of nucleotide diversity, with over 20 million SNPs observed in just six individuals. Consistent with theoretical expectations, we observe reduced diversity in protein-coding sequences, reflecting strong background selection and functional constraints. McDonald-Kreitman tests further reveal that the majority of non-synonymous substitutions between species were driven by positive selection, as expected when selection is highly efficacious in large populations. In addition to providing high-quality genomic resources for a genomically understudied group, our assemblies lay the groundwork for future work in *B. glandula* and other barnacles as models for the study of evolution in large, hyperpolymorphic populations.

## Materials and Methods

### DNA and RNA sequencing

#### High-molecular weight DNA and PacBio sequencing

For genome sequencing using PacBio Hi-Fi reads, we sampled several *Balanus glandula* individuals from the docks in the Oregon Institute of Marine Biology, in Charleston, Oregon, USA (43.346 N, 124.325 W). Individuals were hand collected at low tide in February 2023. High-molecular weight (HMW) DNA was extracted using the PacBio Nanobind tissue kit (PacBio) and quality checked using fluorometric quantification of concentration (Qubit; Invitrogen) and fragment length distribution (Fragment Analyzer; Advanced Analytical). The extracted DNA of a single individual was sequenced on three SMRTcells of a PacBio Sequel II machine at the University of Oregon’s Genomics and Cell Characterization Core Facility (GC3F). These three sequencing runs yielded a total of 5.93 million reads with a read N50 of 12.4 kbp.

#### Chromosome conformation capture

We generated chromosome conformation capture (Hi-C) libraries from a *B. glandula* individual collected from Cape Perpetua, Oregon, USA (44.279 N, 124.112 W) in August 2023. Library construction was performed by Phase Genomics, Inc. The final library was sequenced on an Illumina NovaSeq6000 S4 lane at the University of Oregon’s GC3F, yielding 414.3 million read pairs.

#### ONT ultra-long read sequencing

The HMW DNA from a *B. glandula* individual collected from Cape Perpetua, Oregon, USA in August 2023 was extracted using the QIAGEN Genomic-Tips 20/G kit, processed using an Ultra-Long DNA Sequencing Kit and sequenced using an Oxford Nanopore (ONT) GridION at the University of Oregon’s GC3F. This library yielded 387 thousand ONT reads with a read N50 of 42.7 kbp.

#### RNA sequencing

RNA was extracted from a *B. glandula* individual collected from Cape Perpetua, Oregon, USA in August 2023. For RNAseq, RNA was extracted from the testes and cirri, and sequenced on an Illumina NovaSeq6000 S4 lane at the University of Oregon’s GC3F, resulting in 276.8 million read pairs across both tissue types. The same extracted RNA was used for Iso-Seq, preparing libraries from cirri and testes, and sequenced on a PacBio SMRTcell at the University of Oregon’s GC3F. The testes library yielded 1.4 million reads with a read N50 of 1.4 kbp, while the cirri library yielded 1.9 million reads with a read N50 of 1.5 kbp. In total, the two libraries combined resulted in 3.3 million reads, with a read N50 of 1.5 kbp.

#### Whole-genome shotgun sequencing

Genomic DNA was extracted from eight *B. glandula* individuals collected from Cape Perpetua, Oregon, USA in August 2023 using the QIAGEN Genomic-Tips 20/G kit. The extracted DNA was processed using the Illumina TrueSeq library kit and sequenced, pooled along with other samples, on two Illumina NovaSeq6000 S4 2 *×* 150 bp lanes at the University of Oregon’s GC3F. Across both lanes, the sequencing process yielded 1.97 billion reads across the eight individuals, an average of 246.3 million reads per individual. In total, these individuals were sequenced at an estimated depth of *≈* 60x.

### Genome assembly

#### Genome size and heterozygosity estimation

We estimated the genome size and heterozygosity of the *B. glandula* genome using *k*-mers. First, we used the count command of the jellyfish software version 2.2.10 (Marçais & Kingsford, 2011) to count the unique *k*-mers present in the PacBio HiFi reads. We counted *k*-mers of size 21 (--mer-len 21) present in both strands (--canonical). From these counts, we then generated an empirical *k*-mer distribution using jellyfish’s histo command. Finally, we used GenomeScope2 version 2.0 (Ranallo-Benavidez et al., 2020) to model the distribution of *k*-mers in the diploid genome (--ploidy 2) based on the empirical counts.

#### Base contig-level genome assembly

A base, contig-level genome assembly for *B. glandula* was generated from the PacBio HiFi reads using hifiasm version 0.19.6-r595 (H. Cheng et al., 2021, 2022) with default parameters. This base assembly was assessed for contiguity using quast version 5.2.0 (Gurevich et al., 2013), and for gene-completeness with compleasm version 0.12-r237 (Huang & Li, 2023) using the arthropoda_odb10 reference BUSCO dataset (Manni et al., 2021; Simão et al., 2015; Tegenfeldt et al., 2025) (Table S1).

#### Assembly decontamination

Following the generation of a contig-level genome assembly, we detected exogenous contamination (i.e., sequences not belonging to *B. glandula*) in both reads and assembled contigs using the BlobToolKit software version 4.3.0 (Challis et al., 2020). To achieve this, we first created a new blobtools database, using the create command, specifying the contig-level assembly as our reference sequence (--fasta), the NCBI taxonomic identifier for *B. glandula* (--taxid 110520), and the NCBI reference taxonomy files (--taxdump, downloaded on 2023/12/13).

To add depth of coverage to this database, we aligned our PacBio HiFi sequencing reads to our base, contig-level reference assembly using minimap2 version 2.26-r1175 (H. Li, 2018), running in the map-hifi configuration (-x map-hifi) and exporting alignments in SAM format (-a). Alignments were then processed, sorted, and indexed using samtools version 1.18 (H. Li et al., 2009). The processed alignments were then added to the blobtools database using the add command.

We performed taxonomic assignment of each sequence in the assembly by first aligning our contigs against NCBI’s non-redundant (nr) database (downloaded on 2023/12/13). We aligned each contig against the database using blastn version 2.15.0+, as implemented in BLAST+ (Camacho et al., 2009), specifying a minimum e-value cutoff (-evalue) of 1 *×* 10*^−^*^10^, exporting hits for a maximum of 25 target sequences (-max_target_seqs 25), and exporting results in tabular format (-outfmt 6). These BLAST alignments were then added to the blobtools database using the add command, specifying the rule for taxonomic assignment (--taxrule bestsumorder). Using this approach, the blobtools database provides a single taxonomic assignment to each contig.

Following taxonomic assignment of the assembled contigs, we used blobtools’s filter command to obtain the IDs of sequencing reads that were assigned to a given taxon. We specified paths of our reference contigs (--fasta), alignments (--cov), and sequencing reads (--fastq). For filtering, we specified the parameters to select the reads aligned to contigs assigned to the class Thecos-traca (--Keys=Thecostraca) using a given taxonomic assignment rule (--param bestsumorder_class).

We then used the (--invert) option to specify that we wished to export the IDs of reads matching this assignment (i.e., keeping only reads specific to the barnacle class Thecostraca).

Lastly, we used the IDs of the Thecostraca-assigned reads to filter our raw PacBio HiFi FASTQ file using the subseq command from the seqtk software version 1.4-r130 (H. Li, 2023). This filtering retained 5.74 million Thecostraca-specific PacBio HiFi reads, representing 96.8% of the total sequencing library. This taxonomic assignment and read filtering was also performed on the ONT reads, retaining 371 thousand (95.8%) reads.

#### Optimized contig-level genome assembly

We generated a new contig-level genome assembly using the filtered data and iterating over different assembly parameters and filtering schemes, selecting an optimal assembly based on the best contiguity and gene-completeness metrics (Rayamajhi et al., 2022), as determined by quast and compleasm, respectively. This optimized assembly was generated using hifiasm version 0.19.8-r603, including our filtered PacBio HiFi reads as well as the ultra-long ONT (--ul). This assembly was done using an estimated haploid genome size of 800 Mbp (--hg-size 800m), dropping *k*-mers observed over ten times the average coverage (-D 10.0), and allowing scaffolding (--dual-scaf). For identifying and correctly phasing haplotype sequences, we set the expected coverage at homozygous sites (--hom-cov 68) and the upper bound of coverage to purging duplicates (--purge-max 68) at 68x, along with setting the similarity threshold between duplicate haplotigs to 25% (-s 0.25). This assembly was composed of 2,218 contigs, with a total length of 1.29 Gbp, and a contig N50 of 1.16 Mbp.

Following the initial assembly of contigs, we purged haplotig sequences using the purge_dups software version 1.2.5 (Guan et al., 2020). Following the purge_dups documentation, we first mapped the PacBio HiFi reads to this contig-level assembly using minimap2 -x map-hifi. We then calculated the read-depth histogram using the pbcstat command, followed by determining the base-level coverage cutoffs using the command calcuts. Subsequently, we split the assembly at gaps, if present, using the split_fa command, followed by a self-alignment using minimap2 -x asm5. The purge_dups command was then used to mark haplotig sequences in a BED file according to the coverage cutoffs empirically determined from the data. Lastly, the assembly FASTA was processed to remove sequences tagged as haplotigs using the get_seqs command, only removing sequences at the ends of contigs (-e) and allowing the splitting of contigs (-s). The purging process removed 876 fragments from the primary assembly, accounting for *≈* 240 Mbp of haplotig sequences (Table S1).

#### Hi-C scaffolding

We scaffolded the purged contig-level assembly into chromosome-level scaffolds using Hi-C data. First, we aligned the Hi-C short-read data to our assembly using bwa mem version 0.7.17-r1188 (H. Li, 2013), setting the minimum alignment score to output to zero (-T 0), skipping mate rescue (-S), mate pairing (-P), and setting the smallest query coordinate as primary for split alignments (−5). Alignments were converted to BAM format using samtools version 1.18.

We processed the aligned Hi-C data using pairtools version 1.0.2 (Open2C et al., 2023), following the Dovetail analysis pipeline (Dovetail Genomics, 2023). Briefly, we recorded valid ligation events using pairtools parse, setting a minimum mapping quality of 40 (--min-mapq 40), a maximum inter-alignment gap of 30 kbp (--max-inter-align-gap 30), and reporting the 5’-most unique alignment when detecting multiple ligation events (--walks-policy 5unique). We then sorted the processed alignment pairs using pairtools sort, and marked PCR duplicate reads using pairtools dedup --mark-dups. Finally, we used pairtools split to process the data into a BAM file of valid alignments and a “pairs” file with the read pairs forming the contact map. The resulting alignment BAM was sorted and indexed using samtools.

The processed Hi-C alignments were used to scaffold the genome using the YaHS software version 1.2a.1 (Zhou et al., 2023). Since our assembly consisted of highly fragmented contigs, we reduced the minimum resolution size to 1 kbp (-r 1000). Following the YaHS documentation, we generated a Hi-C contact map using the juicer suite of tools (Dudchenko et al., 2018; Durand et al., 2016). First, we used the juicer pre command (version 1.1) to convert the YaHS output into the juicer format in assembly mode (-a). We then ran juicer tools version 1.9.9 to generate the *.assembly and *.hic files required for manual curation using the Juicebox GUI version 2.17.00. Following manual curation of the contact map, we regenerated the assembly using juicer post. This scaffolded assembly was composed of 650 fragments, a total length of 1.05 Gbp, and a scaffold N50 of 50.96 Mbp (Table S1).

#### Polishing and correcting

After manual curation of the scaffolded genome, we performed one round of polishing and automatic curation using the Inspector software version 1.0.1 (Y. Chen et al., 2021). First, we evaluated the assembly using inspector.py, mapping the PacBio HiFi reads against the assembly. Inspector calculated a mapping rate of 99.98% of these reads to the assembly (87.94% for larger contigs) and a quality value (QV) of 30.49. It additionally identified 742 structural errors larger than 50 bp, including 357 haplotype switches, 303 sequence expansions, and 16 inversions. Assessing smaller (*<*50 bp) errors, Inspector identified 1,929 small-scale expansions, 1,756 small-scale collapses, and 15,618 base substitutions. After evaluation, the assembly was then processed using inspector-correct.py (Table S1) to export the corrected scaffolds in FASTA format.

### Genome annotation

#### Repeat annotation

We identified and classified repetitive elements in the *B. glandula* genome using the EarlGrey software version 4.1.0 (Baril et al., 2024). Before running EarlGrey, we first downloaded the Dfam 3.8 database (Storer et al., 2021), extracting the annotated crustacean repeat sequences to generate an initial consensus masking library. We provided this initial consensus library to EarlGrey (-l). We also used “arthropoda” as the search term for RepeatMasker version 4.1.2 (Smit et al., 2013). In addition, we clustered the final TE library to reduce redundancy (-c yes) and generated a soft-masked genome (-d yes).

Since the homology-based repeat classification from EarlGrey produces a large fraction of unknown elements (*>*35% of the assembly), we ran additional classification steps on this unknown subset of identified repeats. First, we ran TEsorter version 1.5.1 (Zhang et al., 2022), searching against the REXdb version 3.1 metazoan repeat domain database (Neumann et al., 2019) using HMMER version 3.4 (Eddy, 2011). The resulting unknown fraction was then classified using DeepTE (commit 1f76ff9; H. Yan et al., 2020), which uses a convolutional neutral network pre-trained on the *k*-mers of known repeats for TE classification. We ran DeepTE in two rounds. First, we ran DeepTE_domain.py to perform domain assignment on these unknown sequences, using HMMER to search against the supplied minifam database. For the second round, we ran the main classification using DeepTE.py, supplying the found domains to the main classifier (-modify), searching against the pre-trained Metazoan model set (-m Metazoan, -sp M; archived 2019/12/13), and setting a stricter probability threshold of 80% for filtering classified repeats (-prop_thr 0.8).

#### Annotation of protein-coding genes

After masking the repetitive sequences in the genome, we took an integrative approach to annotate protein-coding genes, combining evidence from short-read RNAseq data, PacBio IsoSeq long reads, and protein sequences. For protein-level evidence, we downloaded the Arthropoda and Crustacea partitions of OrthoDB version 11 (Kriventseva et al., 2019; Zdobnov et al., 2021).

When processing the short-read data, we first filtered and processed the raw RNAseq Illumina reads using fastp version 0.23.4 (S. Chen, 2023; S. Chen et al., 2018), setting the minimum length of reads post-filtering to 25 bp (--length_required 25) and enabling the detection of adapter sequences in the paired reads (--detect_adapter_for_pe). We then aligned the processed RNAseq reads to the *B. glandula* assembly using hisat2 version 2.2.1 (D. Kim et al., 2019), exporting alignments tailored for *de novo* transcriptome assemblers (--dta). The alignments were further processed and sorted using samtools.

Using this aligned RNAseq data, we annotated protein-coding genes using BRAKER version 3.0.8 (Gabriel et al., 2024). We ran BRAKER in ETP mode (Brůna et al., 2024), providing both RNAseq alignments (--bams) and Arthropod protein sequences (--prot_seq) as input. We also used the Arthropod dataset from BUSCO (Simão et al., 2015) to enforce a set of orthologs as hints for Augustus (Stanke et al., 2008) and reduce the number of missing single-copy orthologs in the final annotation (--busco_lineage arthropoda_odb10). After running BRAKER, we assessed the quality of the generated annotation using compleasm version 0.2.5 (Huang & Li, 2023) in protein mode, using the Arthropod ortholog set as reference (--lineage arthropoda_odb10).

In parallel, we annotated protein-coding genes using the PacBio IsoSeq data, aligning it to the assembly using minimap2 version 2.26-r1175 (-x splice:hq) and processing the downstream alignments with samtools. We then annotated the genome using the long-read branch of BRAKER version 3.0.8. Similar to the short-read annotation, we ran BRAKER in ETP mode, using the aligned IsoSeq data (--bam) and Arthropod protein sequences (--prot_seq) as evidence, and enforcing the BUSCO Arthropod reference set in the annotation (--busco_lineage arthropoda_odb10). Lastly, we used compleasm protein to assess the completeness of the annotation.

Once the RNAseq and IsoSeq-based BRAKER annotations were completed, we merged the two using TSEBRA (Gabriel et al., 2021). As recommended by published protocols (Brůna et al., 2025), we merged the two annotations by providing the hint files (--hintfiles) and final GTF (--gtf) of each individual BRAKER run as inputs to the tsebra.py script. We assessed the gene-completeness of the merged annotation using compleasm protein, using both the arthropoda_odb10 and crustacea_odb12 reference datasets.

### Transcript and isoform curation

We performed an additional curation of the annotated protein coding sequences using our PacBio Iso-Seq data. First, transcripts and isoforms were annotated from the Iso-Seq long reads using the nf-core/isoseq pipeline version 2.0.0 (Guizard et al., 2023). Quality control of the annotated isoforms was then performed using sqanti3 pipeline version 5.2.2 (Pardo-Palacios et al., 2024). Following the sqanti3 documentation, we first performed quality control of the isoforms using sqanti3_gc, using the BRAKER GTF as our reference annotation, and providing short-read RNAseq reads to infer junction coverage and determine expression levels. Isoforms were then filtered using sqanti3_filter using the default rules, except that we retained mono-exonic transcripts. Lastly, we rescued filtered transcripts and isoforms using sqanti3_rescue, rescuing all isoform types (--mode “full”) and using default parameters.

The rescued isoforms from sqanti3 were processed using TransDecoder version 5.7.1 (Hass, 2023) to identify protein-coding sequences among the annotated sequences. Following the Trans-Decoder documentation, we first extracted the longest coding sequences from the rescued sqanti3 GFF using TransDecoder.LongOrfs. We then identified homology among the putative transcripts using hmmsearch version 3.4 (Eddy, 2011) against the Pfam database (downloaded on 2025/07/25), and using blastp version 2.15.0+ against the UniProtSprot database (downloaded on 2025/08/08). Then, we performed the final coding sequence prediction integrating the homology results using TransDecoder.Predict. Predicted transcripts were mapped back to the genome and to the isoform annotation using the TransDecoder auxiliary scripts, filtering whole transcripts and isoforms lacking homology and coding predictions.

Lastly, we merged this curated isoform annotation with our BRAKER annotation using the tacoRNA software version 0.7.3 (Niknafs et al., 2017). First, we used taco_run to merge the two annotations, disabling the filter by expression levels (--filter-min-expr “0.0”). The merged annotation was then compared to the reference genome and annotation using taco_refcomp to ensure overlap of the merged transcripts. The final annotation was processed using agat version 1.4.3 (Dainat, 2022) to sort by genomic coordinates, add missing annotation features, and standardize annotation IDs. We performed quality control of the merged annotation by assessing gene-completeness using compleasm protein against both the arthropoda_odb10 and crustacea_odb12 reference datasets (Table S2).

### Functional annotation

Functional annotation of the processed protein coding genes was performed using the EnTAP software version 2.3.0 (A. J. Hart et al., 2020). We compared the *B. glandula* protein sequences against the following curated databases: eggNOG (downloaded on 2025/07/08), NCBI nr (downloaded on 2025/10/08), RefSeq Protein (downloaded on 2025/08/08), Swiss-Prot (downloaded on 2025/08/08), and InterProScan (downloaded on 2025/07/25). We considered sequences as contaminant if they had high similarity to bacteria, fungi, viridiplantae, archaea, nematoda, cestoda, and trematoda. Contaminant sequences were removed from the annotation GFF file.

### Annotation of non-coding RNAs

We annotated three classes of non-coding transcripts in the *B. glandula* assembly: ribosomal RNAs (rRNA), transfer RNAs (tRNA), and long, non-coding RNAs (lncRNAs). First, we annotated rRNAs using barrnap version 0.9 (Seemann, 2018), specifying the search against eukaryotic sequences (--kingdom euk). Transfer RNAs were annotated using the tRNAscan-SE software version 2.0.12 (Chan et al., 2021). We specified the annotation of eukaryotic tRNA motifs (-E) and reporting the source of all sequence hits in the outputs (--hitsrc). We removed pseudogenized tRNA copies from downstream analyses.

For annotating lncRNAs, we first re-aligned our PacBio IsoSeq transcripts to the assembly using minimap2 (using the -ax splice:hq option). Then, we called *de novo* transcripts from the aligned long-reads using StringTie version 2.2.1 (Kovaka et al., 2019; Shumate et al., 2022). We ran StringTie in long-read mode (-L), providing our curated protein-coding annotation as a guide for transcript assembly (-G). After calling *de novo* transcripts, we annotated lncRNAs using the FEELnc pipeline version 0.01 (Wucher et al., 2017). In this pipeline, we used FEELnc_filter to filter the StringTie annotation to look for candidate non-coding transcripts not present in the reference protein-coding gene annotation. We then used FEELnc_codpot to calculate the coding potential of the candidate transcripts, modeling the coding potential against a non-coding fraction of the genome (--mode=intergenic). Lastly, we classified candidate non-coding transcripts according to their location relative to nearby genes using FEELnc_classifier.

After their annotation, all three non-coding RNA classes were separately processed using agat in order to sort features by chromosomal coordinates, add missing features, add biotypes, standardize IDs, and merge records into a single GFF file.

### Identifying orthologous gene families

We identified orthologous gene families across annotated barnacle genomes. In addition to the *B. glandula* annotated protein-coding genes, we used annotations for a newly generated assembly and annotation for the wrinkled barnacle (*Balanus crenatus*; see Assembly and annotation of *Balanus crenatus*), as well as published annotations and/or transcriptomic data for the striped barnacle (*Amphibalanus amphitrite*; Han et al., 2024; J.-H. Kim et al., 2019), the Bay barnacle (*Amphibalanus improvisus*; Bishop et al., 2025), and the gooseneck barnacle (*Pollicipes pollicipes*; Bernot et al., 2022). See Processing annotations of published barnacle assemblies for additional details. For all species, we used the longest isoform as representative for each gene for all downstream analyses.

We identified orthologs across the protein-coding genes of these five genomes using OrthoFinder version 3.1.0 (Emms & Kelly, 2015, 2019; Emms et al., 2025). We used the default search parameters in OrthoFinder, inferring gene trees from multiple-sequence alignments (-M msa), searching homologous sequences using diamond version 2.1.9.163, generating multiple-sequence alignments using famsa version 2.4.1, and building gene trees using fasttree version 2.1.11. Additionally, we split paralogous claded below the root into separate orthogroups (-y). Across all genomes, we assigned 88.5% of genes into 15,184 orthogroups, including 2,576 containing single-copy orthologs present in all five species.

### Conserved synteny analysis

We identified regions of conserved synteny between barnacle genomes using the software Synolog version 1.0 (Madrigal & Catchen, 2026). First, we used the synolog_cctl.py script to build a new Synolog cache (new command). The assembly and annotation for the target genomes (*B. glandula*, *A. amphitrite*, and *P. pollicipes*) were then added to this cache using the species, annotation, genes, and integration commands. After the cache was populated, we used the synolog_blastctl.py to run pairwise sequence comparisons between all species, using blastp version 2.15.0. The core Synolog algorithm was then executed using the synolog_run.py script. Genome-wide synteny plots were generated using the synolog_plot.py genome command, plotting only orthologs identified in all species (--span), coloring orthologs by their chromosome of origin (--color 0), and only plotting sequences larger than 5 Mbp (--min 5e6).

### Identifying highly-conserved elements

We identified highly-conserved genomic elements (HCE) across the genomes of six barnacle species (*B. glandula*, *B. crenatus*, *A. amphitrite*, *A. improvisus*, *C. mitella*, and *P. pollicipes*). First, we generated a whole-genome alignment of these five species using cactus version 2.9.7 (Armstrong et al., 2020). We used the phylogenetic tree generated by OrthoFinder as the guide tree for the alignment. Next, we ran the phyloFit program, part of phast version 1.5 (Siepel et al., 2005) to generate a neutral phylogenetic model based on the resulting alignment and general time reversible substitution model (--subst-mod REV). Lastly, we calculated conservation scores on the aligned sequences using phastCons. We exported the coordinates (--most-conserved) and assigned log-odd scores (--scores) to the HCEs.

We used a custom Python script (tally_phastcon_elements.py) to annotate and calculate the enrichment of known annotated features present in the phastCons. The script provides general statistics about the HCEs (e.g., number of HCE and size distribution) and annotates conserved sites based on their overlap to annotated features in the genome.

### Estimating rates of non-synonymous to synonymous substitution

We calculated pairwise d_N_/d_S_ between the coding sequences of *B. glandula* and its congener *B. crenatus*. We first took the orthologous protein coding genes identified between the two species by OrthoFinder. Then, we used a custom Python script (extract_orthogroups_cds.py) to extract the per-species coding sequences for each single-copy ortholog between the two species. We generated a multiple-sequence alignment (MSA) for each ortholog using prank version v.170427 (Löytynoja & Goldman, 2005) in codon mode (-codon), using the OrthoFinder phylogeny as the guide tree for alignment (-tree; -prunetree), and performing three iterations of re-alignments (-iterate=3). The MSAs were then processed using ClipKIT version 2.7.0 (Steenwyk et al., 2020), trimming the alignment by codons (--codon). Lastly, we filtered all alignments shorter than 100 bp.

After generating and processing alignments for the resulting 9,082 single-copy orthologs, we calculated d_N_/d_S_ rates using a custom Python script (pairwise_dnds.py). The script uses the dnds Python package version 2.1 (Qalieh, 2023) to calculate d_N_/d_S_ using the codon-by-codon counting method described by Nei and Gojobori (1986). For each alignment, we calculated the total number of non-synonymous and synonymous sites (N_C_ and S_C_, respectively), accounting for the degenerate codons. We then calculated the number of non-synonymous (N_D_) and synonymous differences (S_D_) observed between the two sequences. The rate between the number non-synonymous (d_N_) and synonymous (d_S_) substitutions and sites is then calculated, and corrected for multiple substitutions using the Jukes-Cantor correction (Jukes & Cantor, 1969).

### McDonald-Kreitman test

We calculated the deviation from synonymous to non-synonymous polymorphism and divergence using the McDonald-Kreitman (MK) test (McDonald & Kreitman, 1991), using the single-copy orthologs between *B. glandula* and *B. crenatus* described above. For determining the levels of polymorphism, we took population genetic data from *B. glandula* from Central Oregon (see Population genetics of central Oregon barnacles). We used a custom Python script (extract_hap_cds.py) to map these variants (both SNPs and indels) to the *B. glandula* coding sequences, effectively creating two allelic variants for each sample for each protein coding gene. This polymorphism data was re-organized for each ortholog, generating a per-ortholog multi-species FASTA file, containing both polymorphic sequences for *B. glandula* and the corresponding *B. crenatus* outgroup sequence. Similar to the d_N_/d_S_ analysis, we generated codon-aware multiple-sequence alignments of all coding sequences using prank -codon. Orthologs whose alignments contained frameshift mutations were removed from the analysis. The resulting alignments were processed with ClipKIT, trimming the alignment by codons (--codon), and trimming sites above a gap threshold of 25% (--mode gappy; --gaps 0.25). Alignments smaller than 100 bp were also discarded, retaining sequences for 7,374 orthologs.

The MK test and associated statistics were calculated using the MKado version 0.2.0 software (Rivera-Colón et al., 2026). MKado parses the multiple-sequence alignments and computes the non-synonymous and synonymous changes in divergence and polymorphism (D_N_, D_S_, P_N_, P_S_, respectively), between ingroup and outgroup sequences. We ran MKado three times, each time calculating a different variant of the MK test. First, we calculated the standard MK test (McDonald & Kreitman, 1991) for all 7,374 orthologs using the MKado batch command, calculating the pergene proportion of non-synonymous sites fixed by selection (*α*; N. G. C. Smith and Eyre-Walker, 2002), Neutrality Index (NI; Rand and Kann, 1996) and Direction of Selection (DoS; Stoletzki and Eyre-Walker, 2011). Next, we calculated the asymptotic MK test (--asymptotic; Messer and Petrov, 2013) to correct for the biases introduced by low frequency polymorphism, and estimating *α*_asym_ aggregated across all genes. Lastly, we calculated the Tarone-Greenland weighted estimator (*α*_TG_; Stoletzki and Eyre-Walker, 2011) using MKado’s --alpha-tg, which provides an unbiased estimator of genome-wide *α* when comparing across multiple genes.

### Gene ontology enrichment

We calculated functional enrichment among our candidate genes under positive selection using the topGO R package version 2.62.0 (Alexa & Rahnenführer, 2017), along with the Gene Ontology (GO) annotation available in GO.db version 3.22.0 (M. Carlson, 2017; released on 2025/07/22). First, we took the 7,374 genes analysed with MKado’s MK test and linked them to the functional annotation (including GO terms) generated by EnTAP. In total, we retained 5,101 genes having both MK test results and annotated GO terms. This gene subset included 367 of our candidate genes under positive selection. Then, for each of the three GO domains (Biological Process, Cellular Component, and Molecular Function), we used the annFUN.gene2GO function to build a topGOdata object mapping genes and GO terms, pruning terms with fewer than five annotated genes. Using the runTest function we tested for GO term enrichment in our 367 candidate gene set against the 5,101-gene background using a Fisher’s exact test. In addition to the default parameters, we ran this test with the elim topology-aware algorithm, which accounts for the hierarchical nature of GO terms to prevent the inflation in a term’s enrichment originating from parent-child relationships in the GO graph (Alexa et al., 2006), i.e., genes annotated to a significant child term are removed before its parent terms are tested. Per-term counts and *p*-values were then extracted using the GenTable and score functions.

### Codon usage biases

We calculated codon usage biases on the *B. glandula* coding sequences using the R package cubar version 1.2.0 (M. Liu et al., 2026). Before running cubar, we processed the annotation to extract only the longest isoform for each gene using agat’s agat_sp_keep_longest_isoform.pl and agat_sp_extract_sequences.pl. In cubar, we first filtered coding sequences using the check_cds function, ensuring the presence of canonical start and stop codons. To prevent known biases in small coding sequences, we then removed transcripts smaller than 300 bp and those above and below two standard deviations of the mean transcript length. This process retained 16,217 sequences in *B. glandula*. After obtaining the empirical codon distribution using the count_codons function, we then used get_enc to calculate codon usage biases using the effective number of codons (ENC) metric, which captures the deviation from equal codon usage across synonymous codons (Wright, 1990). More specifically, we calculated an improved implementation of ENC, accounting for usage differences across different codon subfamilies (Sun et al., 2013). In addition to ENC, we calculated GC proportions using the following cubar functions: get_gc for coding sequence-wide GC proportion, get_gc3s for GC proportion at 3rd codon sites (GC3s), and get_gc4d for GC proportion at 4-fold degenerate sites (GC4d). Lastly, we used Wright’s curve (ENC*_exp_* = 2 + *s* + 29*/*[*s*^2^ +(1 *−s*)^2^], were *s* denotes GC3s; Wright, 1990) to determine the expected distribution of the number of effective codons given a given GC3s distribution. At each gene, we calculated its deviation from this empirical ENC distribution using ENC*_dev_* = (ENC*_exp_ −* ENC*_obs_*)*/*ENC*_exp_*, as well as a linear regression of ENC*_obs_* given ENC*_exp_* using R’s lm function. We compared the ENC results in *B. glandula* by calculating empirical and expected ENC distributions for 11 other metazoan genomes, as specified in Table S6.

### Genetic variation in the Balanus glandula reference

We genotyped our barnacle reference individual to assess the levels of genetic variation present in our genome assembly. To achieve this, we first re-aligned our processed PacBio HiFi reads to the assembly using minimap2 version 2.28-r1209 with the -x map-hifi preset. A total of 99.97% of reads were successfully mapped to the reference. The aligned reads were then sorted and filtered using samtools version 1.21 to remove non-primary and supplementary alignments. Lastly, we used the mosdepth software version 0.3.10 (Pedersen & Quinlan, 2018) to calculate the depth of coverage along 10 kbp windows (--no-per-base; --by 10000) and calculating the proportion of bases covered at specified coverage thresholds (--thresholds “1,3,5,10,20”). After filtering and processing alignments, the reference individual retained a mean depth of coverage of 50.88x (median: 47.94x, standard deviation: 100.32x).

Genotypes were called from these alignments using the standard bcftools version 1.21 (Danecek et al., 2021) genotyping pipeline, consisting of the mpileup and call commands for generating genotype likelihoods and calling genotypes, respectively. We only genotyped the 58 contigs larger than 1 Mbp, processing each contig separately (using --regions). For mpileup, we only used a maximum depth of 250 x (--max-depth 250), and adding an annotation for both per-sample and per-site total and allelic depth (--annotate “FORMAT/AD,FORMAT/DP,INFO/AD”). For the call command, we used the updated calling model (--multiallelic-caller), adding annotations for genotype qualities and posterior probabilities (--annotate “GQ,GP”). Additionally, we used the bcftools norm command to join adjacent SNPs and indels into single multiallelic sites.

After calling genotypes, we filtered the resulting variants using bcftools. Using the filter command, we removed sites (both variant and invariant) with low quality (--exclude “QUAL < 30”), with too high coverage (--exclude “INFO/DP > 125”), adjacent to indels (--SnpGap 3; --IndelGap 5), and within the span of annotated repeats (--mask-file repeats.bed). We fil-tered individual genotypes with a low genotyping quality (--exclude “FORMAT/GQ < 30”), low depth (--exclude “FORMAT/DP < 10”), and a large allelic imbalance (--exclude “(GT==’het’) & %MAX(FMT/AD)/(FMT/DP) < 0.3”). We also removed sites with more than two alleles using (--max-alleles 2) from the view command. Lastly, we merged all the files together in a single genome-wide VCF using bcftools concat.

We classified the effects of heterozygous variants present in the reference genome using snpEff version 5.2 (Cingolani et al., 2012). We first built a snpEff database of our *B. glandula* reference genome and annotation using the build command. Then, we estimated the effects of variants using the eff command, exporting a CSV file containing a breakdown of all annotated variants (-csvStats).

### Population genetics of central Oregon barnacles

#### Aligning reads

We used whole-genome shotgun sequencing in order to assess the genetic diversity of a *B. glandula* population in central Oregon. We aligned the Illumina paired-end reads to the reference genome using bwa mem version 0.7.18-r1243. Resulting alignments were processed and sorted using samtools version 1.21. We used Picard MarkDuplicates version 3.3.0 (Broad Institute, 2024) to identify PCR duplicate reads, and gtak LeftAlignIndels version 4.6.1.0 (DePristo et al., 2011; McKenna et al., 2010) to process indels. Then, we used samtools view to filter and export the alignments, removing those with low mapping quality (-q 30), unmapped reads (-F 4), non-primary (-F 256) and secondary (-F 2048) alignments, and those flagged as PCR duplicates (-F 1024). Lastly, we calculated depth of coverage using mosdepth. Two individuals had low genome-wide average coverage (*<*5x) and were removed from downstream analysis. For the six remaining individuals, an average of 94.13% of reads aligned to the reference genome, resulting in a mean genome-wide coverage of 48.83x.

#### Genotyping

These six individuals were genotyped using the same bcftools mpileup/call approach used for our reference sample (see Genetic variation in the *Balanus glandula* reference), except allowing for a maximum depth of 1000x when genotyping (--max-depth 1000). After normalizing indels using bcftools norm, we filtered the called variants using bcftools filter and view. We removed sites with low quality, adjacent to indels, and within the span of annotated repeats. Additionally, we removed sites with per-sample coverage lower than 10x, genotype quality lower than 30, and allelic imbalance larger than 30%. After merging individual per-chromosome variants into a single genome-wide VCF, we also removed any sites containing missing genotypes (bcftools view -g ^miss). The effects of the kept variants was determined using snpEff.

#### Population genetic statistics

We used the pixy software version 2.0.0.beta12 (Bailey et al., 2025; Korunes & Samuk, 2021) to obtain genome-wide measures of genetic diversity. We calculated both nucleotide diversity (*π*) and Watterson’s Theta (*θ*_W_; --stats pi watterson_theta) on 10 kbp windows (--window_size 10000). We allowed multiallelic sites to be used in the calculations (--include_multiallelic_snps), limiting the calculation to sites present in our input VCF file (--sites_file).

We performed additional diversity calculations with pixy on different regions of interest in the genome. Instead of running on uniform 10 kbp windows along the genome, we calculated *π* along the span of annotated genomic features (coding sequences, introns, 5’ UTRs, 3’ UTRs, lncRNAs, rRNAs, and intergenic regions). We reran pixy separately for each feature, specifying the genomic coordinates of the feature’s annotation using the --bed_file option. We also calculated *π* at 0-fold and 4-fold degenerate sites. For this, we first determined the degeneracy at each coding site in the genome using the degenotate version 1.3 tool from the snpArcher pipeline (Mirchandani et al., 2023). After determining degeneracy we separately calculated *π* for 0-fold and 4-fold degenerate sites using pixy, specifying the specific sites of interest using --sites_file and averaging within the span of each exon (using --bed_file).

#### Inference of effective population size

We used our filtered whole-genome genotypes from central Oregon *B. glandula* to estimate the population’s effective size. First, we estimated the long-term average N_e_ in this population based on the estimates of *θ*_W_ along 10 kbp windows using pixy, and using the relationship N_e_ = *θ*_W_*/*4*µ*. We used the per-base per-generation mutation rate (*µ*) of 3 *×* 10*^−^*^9^ estimated by Nunez et al. (2020) for the northern acorn barnacle *Semibalanus balanoides*. Using this relationship, we calculated N_e_ along these 10 kbp windows, obtaining a distribution of effective size along the genome.

Using the same set of filtered genotypes, we additionally estimated the coalescent-based effective size of the population using the msmc2 software version 2.1.4 (Schiffels & Wang, 2020), as implemented in the DemInfHelper program version 0.1.1 (Forest et al., 2024). We ran msmc2 with 25 iterations (-i 25), using the default time segments (-p 1*2+25*1+1*2+1*3), a generation time of 1 (to keep the inference in generations instead of years), and a *µ* of 3 *×* 10*^−^*^9^. We estimated historical N_e_ for our six Cape Perpetua individuals; however since our genotypes were not phased, we ran msmc2 for each individual separately.

## Results

### Genome assembly and annotation

We assembled a chromosome-level reference genome for the Pacific acorn barnacle using PacBio HiFi long reads and Hi-C data. Preliminary *k*-mer assessment of the HiFi reads revealed an expected genome size of *≈* 845 Mbp, with a heterozygosity of over 3% (Figure S1). After purging haplotigs, scaffolding, and polishing the sequences (see Genome assembly), the curated assembly was composed of 592 fragments, and had a total sequence length of 1.04 Gbp and a scaffold N50 of 51.41 Mbp (Table S1). The assembly was 94.08% gene-complete, with 4.64% duplicates, for the arthropoda_odb10 BUSCO dataset (Manni et al., 2021; Simão et al., 2015; Tegenfeldt et al., 2025). Of the 1.04 Gbp assembled, 906.6 Mbp (86.74%) was contained in 58 scaffolds larger than 1 Mbp. Moreover, the 16 largest fragments (*>* 8.5 Mbp; Figure S2) contained 841.2 Mbp (80.61%) of the total sequence, and likely represent 16 chromosomes. This karyotype number, N=16, is in agreement with that observed for other species in the genus *Balanus* (Austin et al., 1958) and other acorn barnacles in the family Balanidae (Han et al., 2024).

Following assembly, we annotated and masked repeat sequences in the genome using EarlGrey (Baril et al., 2024). This analysis identified 687.9 Mbp (65.92%) of the assembly as repetitive elements; however, the majority of these, 404.5 Mbp (38.76% of the assembly), were unable to be classified among the major repeat families. This large fraction of unclassified repeats is likely the product of the small number of representative barnacle sequences among reference repeat databases, further reflecting the status of barnacles as a genomically understudied group. Re-classification of this unknown fraction by TEsorter (Zhang et al., 2022) and DeepTE (H. Yan et al., 2020) reduced the fraction of unclassified elements to only 5.89% of the total assembly. This classification approach identified DNA repeat elements as the most common repeat class in the *B. glandula* genome, comprising 36.3% of the total assembly (Table S3). LINEs and LTRs were the second and third most common repeat types, comprising 10.26% and 4.81% of the assembly, respectively.

We annotated protein-coding genes in the *B. glandula* assembly using a combination of short-read RNAseq and long-read PacBio Iso-Seq data. This process resulted in 20,234 protein-coding genes annotated, which had a BUSCO completeness of 93.58% and 84.77% for the arthropoda_odb10 and crustacea_odb12 datasets, respectively (Table S2). Comparing this annotation against the proteomes of four other barnacle species assigned orthology to 16,978 (83.91%) *B. glandula* protein-coding genes, including 2,576 single-copy orthologs across all five species. In addition to protein-coding genes, we also annotated non-coding elements in the genome, identifying 7,532 long, non-coding RNAs (lncRNA), 1,203 transfer RNAs (tRNA), and 24 ribosomal RNA (rRNA) genes. In total, across protein-coding and non-coding transcripts, we were able to annotate 28,993 genes in the *B. glandula* reference assembly.

Together, the high contiguity, gene-completeness, chromosome-level scaffolding, and comprehensive annotation of this assembly make it among the most complete genomic resources available for any barnacle species, and a strong foundation for downstream population genomic and comparative analyses.

### Conserved architecture across barnacle genomes

To assess large-scale structural conservation across barnacle genomes, we performed a conserved synteny analysis using Synolog (Madrigal & Catchen, 2026). We compared our *B. glandula* genome against the chromosome-level assemblies of the striped barnacle *Amphibalanus amphitrite* (Balanidae; Han et al., 2024) and the gooseneck barnacle *Pollicipes pollicipes* (Pollicipedidae; Bernot et al., 2022). These genomes display conservation in their large-scale organization, including one-to-one correspondence between 16 putatively orthologous chromosomes across the three species (Figure 1). Smaller deviations from this one-to-one relationship likely represent assembly errors are not reflective to changes in chromosomal evolution in these species. Observing this strong correspondance shows that our scaffolding of the *B. glandula* assembly likely reflects the biological structure of the genome and is not the product of technical artifacts, and further suggests conservation of genome architecture across barnacles spanning *≈* 165 Mya of evolution (Bernot et al., 2022; Han et al., 2024). This striking chromosomal stability provides a shared framework for comparative genomic analyses across the order and suggests that large-scale genome rearrangements have not been a major driver of diversification in barnacles.

**Figure 1:**
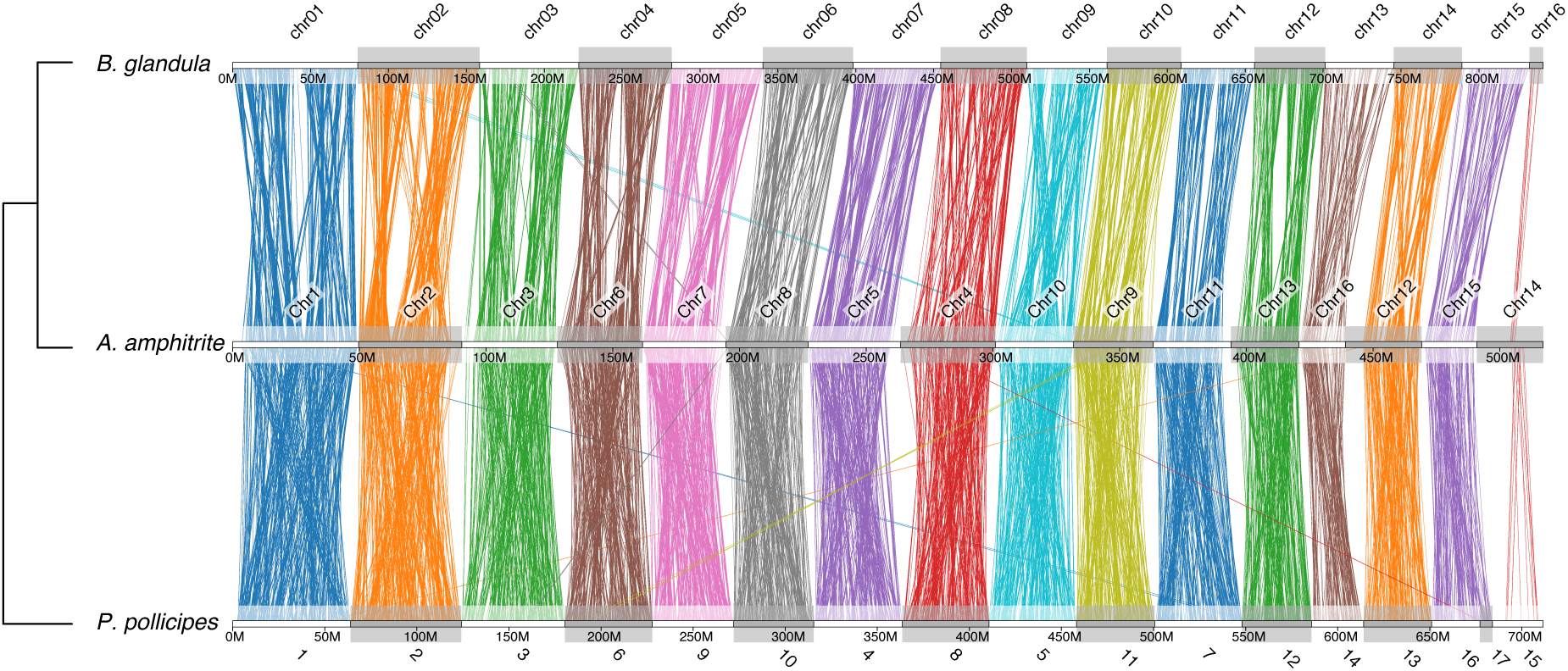
Conserved synteny across barnacle genomes. Genome-wide synteny plots showing one-to-one correspondence across 16 putatively orthologous chromosomes between the genomes of *Balanus glandula* (top), *Amphibalanus amphitrite* (middle), and *Pollicipes pollicipes* (bottom). Each colored line shows protein-coding orthologs across the three assemblies, colored according to their chromosome of origin in *B. glandula*. Cladogram on the left shows the evolutionary relationship among the three species.

### High genetic diversity in the *Balanus glandula* reference assembly

We estimated the genetic diversity present in the *B. glandula* genome by identifying heterozygous variants in our reference individual. After filtering low-quality sites and masking regions with annotated repeats, we obtained genotypes for 314 million sites along the genome, equivalent to 34.71% of the 904.94 Mbp present in the 58 scaffolds larger than 1 Mbp (Table S4). Among these sites, 9.08 million (2.89%) were heterozygous in the reference individual, including 8.73 million SNPs and 356 thousand indels. In total, this variation represents a mean density of 14.66 heterozygous sites per kbp (median: 6 per kbp; standard deviation: 19.86). Moreover, we observed nearly 400 thousand variant sites within protein-coding sequences, including 144 thousand non-synonymous point mutations. Transcripts in the reference individual had a mean of 7.74 (median: 3, standard deviation: 14.97) non-synonymous heterozygous variants, indicating considerable coding variation segregating within a single individual. Even from this single genome, the sheer volume of heterozygosity underscores the exceptionally large population sizes of this species and foreshadows the extreme diversity observed at the population level.

### Functional elements are conserved in barnacle genomes

We used the phastCons package (Siepel et al., 2005) on the cactus whole-genome alignment (Armstrong et al., 2020) between six barnacle species to identify highly conserved elements (HCE) in the Pacific acorn barnacle genome. Across the 16 putative chromosomes in the *B. glandula* assembly, phastCons identified 196,141 HCEs, spanning 50.85 Mbp (6.05%) of the 16 assembled chromosomes. In *B. glandula*, the HCEs had a mean length of 259.25 bp (median: 42 bp, standard deviation: 990.40 bp).

Following the identification of HCEs, we compared their sequence composition against the genome-wide background (Figure 2). We first intersected the genomic coordinates of the HCEs with the *B. glandula* annotation, obtaining the proportion of annotated features among HCE sites (Figure 2A). Among our target features — which include an “intergenic” class for the unannotated fraction of the genome and a “multiple” class for overlapping features (e.g., a site spanning both a transposable element and an intron) — all but one are present among HCEs. We did not observe any conserved rRNA sites; however, this is likely the product of the whole-genome alignment, given rRNA’s repetitive nature and alignment difficulty (J. J. Gillespie, 2004), and not an accurate measure of conservation (or lack thereof). Aside from the absence of rRNAs, the sequence composition of HCEs largely followed that of the whole genome. Most annotated features do not appear substantially enriched among HCE sites (Figure 2B; Table S5). The two clear exceptions are coding sequences (CDSs) and tRNAs. Protein-coding sequences display an approximately 2-fold enrichment among HCEs (mean log_2_ enrichment: 0.75, median: 0.98, standard deviation: 0.93), reflecting the conservation of protein-coding genes among barnacle genomes. Similarly, tRNAs exhibit *≈*4-fold enrichment in HCEs (mean log_2_ enrichment: 2.14, median: 2.04, standard deviation: 0.96), likely reflecting the strong functional constraints on these sequences.

**Figure 2:**
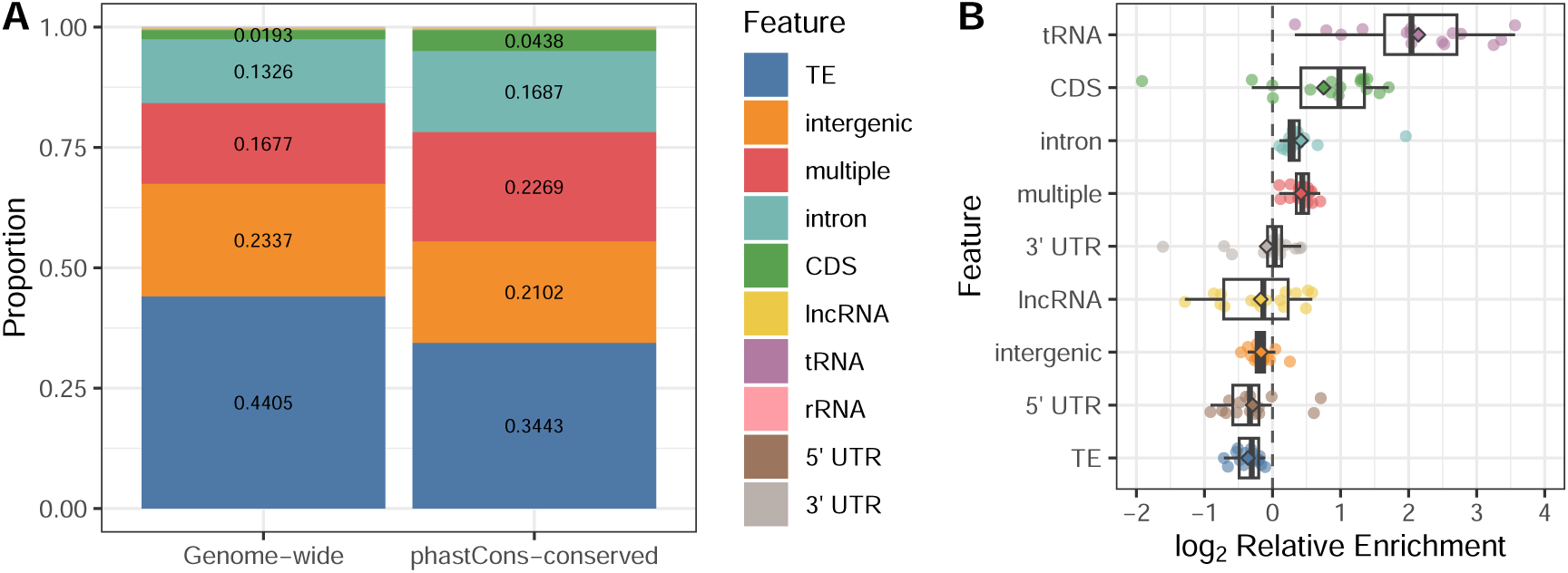
Highly conserved elements in the *Balanus glandula* assembly. **A.** Proportion of annotated features in the whole genome (left column) and the 50.85 Mbp spanning phastCons-conserved elements (right column). Values are shown only for elements with proportions larger than 0.01. Sites that overlap more than one non-intergenic annotation type are represented by the “multiple” feature. **B.** Enrichment of the annotated features in the highly conserved elements compared to the genome-wide background, calculated as the log_2_ of the relative enrichment. The log_2_ enrichment values for each of the 16 chromosomes are shown as points. The diamonds in the boxplot indicate the mean enrichment across the genome.

The enrichment among conserved elements in *B. glandula* mirrors patterns observed across other groups of organisms. First, coding sequences are consistently enriched among conserved elements, following the patterns characterized by Siepel et al. (2005) across vertebrate, insect, worm, and yeast genomes. While the raw composition of enriched conserved elements is different across all these species, the consistent enrichment of CDSs is in accordance with the expectation that protein-coding sequences are among the most functionally constrained genomic regions. Secondly, the enrichment pattern in barnacles follows the same ordering observed across both plants (Hupalo & Kern, 2013) and animals (Siepel et al., 2005): tRNAs are the most enriched annotation among conserved regions, followed by CDSs, while transposable elements are drastically underrepresented among HCEs. In contrast, introns are mildly enriched among barnacle HCEs (Figure 2B; Table S5), while depleted in plant (Hupalo & Kern, 2013), insect, and vertebrate conserved elements (Siepel et al., 2005). However, it remains unknown if this enrichment reflects the larger intronic fraction of the barnacle genome or genuine intronic conservation in this lineage. Overall, the conservation of functional elements across barnacle genomes indicates that purifying selection has been effective at maintaining functionally constrained sequences despite the highly repetitive nature of the genome, a pattern we revisit in the context of the broader selective regime acting on these large populations.

### Patterns of molecular evolution reflect positive selection in protein-coding genes

To examine the rate of molecular evolution in protein-coding sequences, we calculated the pairwise ratio of non-synonymous to synonymous substitutions (d_N_/d_S_) between *B. glandula* and its congener, the wrinkled barnacle *Balanus crenatus*, using the codon counting approach described by Nei and Gojobori (1986). We identified and aligned 9,082 single-copy orthologs between the two species. The mean pairwise d_N_/d_S_ was 0.206 (median: 0.158, standard deviation: 0.191; Figure S3). This ratio indicates that, on average, coding sequences in these barnacle genomes are under purifying selection, reflecting the expected evolutionary constraints on sequences of functional importance.

Of the nine thousand genes assessed, 70 (0.77%) displayed a d_N_/d_S_ *>* 1, and are likely candidates for positive selection. The predicted function of the majority (*>* 80%) of these genes is unknown; however, this is expected, since homology is likely harder to determine for fast-evolving sequences putatively under strong selection. Despite this limitation, we found several genes with known function among our positive selection candidates, including a follistatin-A-like homolog, which is involved in myogenesis and muscle development in crustaceans (Y. Yan et al., 2021). We also observed a homolog of *γ*-interferon-inducible lysosomal thiol reductase, which plays a role in crustacean innate immune response (M. Liu et al., 2019). Additionally, we found evidence of positive selection in homologs of tumor protein p53-inducible protein 11, which is involved in cell cycle regulation and is up-regulated in crabs after viral infection (C.-H. Cheng et al., 2021), as well as a Quiver homolog, a gene related to neuronal activity and sleep regulation in *Drosophila* (Wang & Wu, 2010).

We expanded on our d_N_/d_S_ measures of positive selection in protein-coding sequences in the *B. glandula* genome by incorporating population-level polymorphism using the McDonald-Kreitman (MK) test (McDonald & Kreitman, 1991). Specifically, we used polymorphism data derived from a population panel of six *B. glandula* individuals from the central Oregon coast (see Barnacle populations exhibit elevated genetic polymorphism), comparing those against the orthologous coding sequences in the *B. crenatus* outgroup. After removing alignments shorter than 100 bp and those containing premature stop codons and/or frameshift mutations, we calculated the MK test for 7,374 orthologous sequences between *B. glandula* and *B. crenatus*.

Four hundred and forty-four (444, 6.02%) of these orthologs showed a significant deviation from the expected proportion of non-synonymous and synonymous substitutions and polymorphisms (Figure 3A). We additionally calculated the Neutrality Index (NI; Rand and Kann, 1996) and Direction of Selection (DoS; Stoletzki and Eyre-Walker, 2011) at each ortholog. We observed a mean NI value of 0.745 (median: 0.554; standard deviation: 1.079) and a mean DoS of 0.102 (median: 0.106; standard deviation: 0.131; Figure S4). Both of these statistics (NI *<* 1 and DoS *>* 0) show that on average, and in contrast with our measurements of d_N_/d_S_ (see Molecular evolution in large populations: reconciling d_N_/d_S_ and the McDonald-Kreitman test), protein-coding sequences in *B. glandula* exhibit an excess of non-synonymous substitutions, suggesting that they are evolving under positive directional selection when compared to the outgroup sequence. Moreover, the near absence of genes displaying a significant excess in non-synonymous polymorphism (NI *>* 1) reflect a reduction the proportion of slightly deleterious variants segregating in the population, suggesting the presence of highly efficacious background selection.

**Figure 3:**
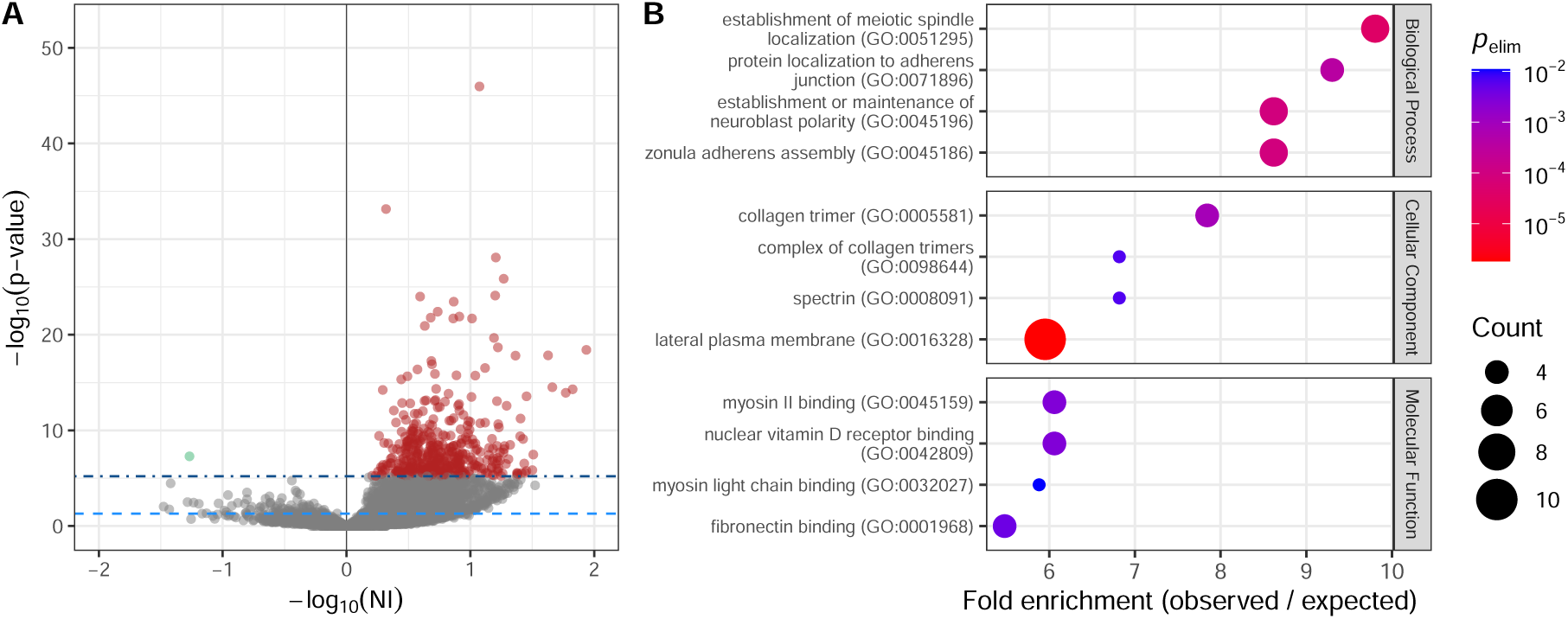
Signatures of positive selection in *Balanus glandula* protein-coding genes. **A.** Selection on 7,374 protein-coding genes was assessed using the McDonald-Kreitman test. X-axis denotes the *−* log_10_ of the Neutrality Index, with values on the right (NI *<* 1) indicating excess of non-synonymous substitutions indicative of positive selection. Y-axis denotes the *−* log_10_ *p*-value of the McDonald-Kreitman test. The dashed and dot-dashed lines show the raw (*p <* 0.05) and Bonferroni-corrected (*p <* 6.20*×*10*^−^*^6^) significance thresholds, respectively. Points in red show the 444 candidates under positive selection, genes with a significant excess of fixed non-synonymous substitutions. Point in green show the one significant gene with an excess of non-synonymous polymorphism. **B.** Significant GO term enrichment among the candidates under positive selection in the McDonald-Kreitman test. *p*_elim_ denotes the *p*-value calculated from topGO’s elim algorithm (Alexa et al., 2006), which accounts for redundancy among GO terms. Top, middle, and bottom panels show the top four terms with the largest enrichment among the candidates under selection for the three GO categories: biological processes, cellular component, and molecular function.

In addition to NI and DoS, we calculated *α*, a measure of the proportion of non-synonymous substitutions fixed by positive selection (N. G. C. Smith & Eyre-Walker, 2002). We calculated two adjusted versions of *α* to account for the presence of known biases. First, we calculated the Tarone-Greenland *α* estimator (*α*_TG_), a weighted estimator that corrects for the effects of averaging over a large number of genes (Stoletzki & Eyre-Walker, 2011). Across our 7,374 genes, we calculated an *α*_TG_ of 0.540 (95% CI: 0.534–0.550), which indicates that *≈* 54% of non-synonymous substitutions in this *B. glandula* population were fixed as a product of positive selection. Secondly, we calculated *α* using the asymptotic MK test (*α*_asym_), which corrects for the presence of weakly deleterious non-synonymous polymorphisms present at low frequency in the population (Messer & Petrov, 2013) which might obscure the signal of positive selection. We found a mean *α*_asym_ of 0.610 (95% CI: 0.610–0.622), a larger proportion than that observed in *Drosophila* (*α*_asym_ = 0.57) and humans (*α*_asym_ = 0.13; Haller and Messer, 2017; Messer and Petrov, 2013). Regardless of the metric used, both *α*_TG_ and *α*_asym_ agree that the majority of non-synonymous substitutions in this barnacle population are the outcome of directional positive selection. Notably, this pervasive signal of positive selection stands in apparent tension with the low genome-wide d_N_/d_S_, a discordance we address in the Discussion.

We explored the predicted function of the candidate genes under positive selection identified in the MK test by performing a gene ontology (GO) enrichment analysis on the 367 candidates with available functional annotation. The most enriched GO terms describe a coherent set of cellular functions, spanning the plasma membrane, adherens junctions, myosin and spectrin complexes, and collagen trimers (Figure 3B; Figure S5). Among the collagen-related terms, we additionally observe the enrichment of processes related to the molting of arthropod cuticles (e.g., molting cycle, collagen and cuticulin-based cuticle). Together, these enrichment patterns suggest strong directional selection in *B. glandula* on genes involved in the building and remodeling of the cell surface, interactions with the extracellular matrix, cell-cell adhesion, and related functions.

Several of our candidates under strong positive selection are direct components of these enriched functions. For example, two of our top candidates are non-muscle myosin heavy chain and myosin-VIIa homologs, both showing a very large fraction of non-synonymous substitutions fixed by positive selection, *α* of 0.961 and 0.940, respectively. There is additional evidence of positive selection on proteins with known direct interactions, such as the components of the spectrin heterodimer, spectrin *β* chain (*α* = 0.898) and spectrin *α* chain (*α* = 0.895), the integrin heterodimer, integrin *α* (*α* = 0.793) and integrin *β* (*α* = 0.810), as well as non-muscle myosin heavy chain (*α* = 0.961; seen above) and its direct activator Rho-associated protein kinase 1 (ROCK1; *α* = 0.939). Observing evidence of positive selection on interacting pairs is strongly suggestive of the coevolution of these sequences, in a compensatory or correlated fashion; however, future research is required to fully characterize the molecular interactions produced by these non-synonymous substitutions in *B. glandula*.

Outside of this cell remodeling and adhesion core, the remaining candidates of selection appear largely functionally heterogeneous, spanning protein synthesis, osmoregulation, and development (e.g., homologs of eukaryotic translation initiation factor 3 subunits L and B, transmembrane channel-like 7, and patched domain-containing protein 3-like, all with *α >* 0.9; Bolatto et al., 2015; Lee et al., 2015; H. Liu et al., 2025). Of these candidates, only a single significant gene in the MK test also appears among our candidates of selection in the d_N_/d_S_ analysis — a putative homolog of formin-like protein 3, a member of a family of proteins known to play a role in crustacean immunity (Y. Li et al., 2025) — further highlighting the discrepancy between the two tests of selection. Finally, our single candidate showing a significant excess of polymorphism relative to divergence is a homolog of tubulin *α*-1C chain-like, a major constituent of microtubules and thus directly linked to the cell-remodeling machinery under positive selection. The large excess of polymorphism at this gene (NI = 18.54) instead suggests balancing selection, and this homolog has additionally been observed as differentially expressed in response to osmotic stress in barnacles (Chandramouli et al., 2015).

### Barnacle genes show biases in the use of synonymous codons

We compared biases in the usage and composition of synonymous codons in 12 metazoan genomes, including four barnacles (Table S6), to study the effects of translational selection in the context of large populations. Using the annotated transcript sequences in these genomes we calculated the effective number of codons (ENC), a measure of the extent of departure from equal usage across synonymous codons (Sun et al., 2013; Wright, 1990). In addition to ENC, we also estimated compositional biases by calculating both the overall coding sequence GC proportion, and the proportion of GC bases at third codon positions (GC3s). Across 16,217 genes in *B. glandula*, we find a mean ENC of 43.17 (median: 42.84, standard deviation: 5.52; Figure 4A; Table S6). This number of effective codons is similar to the one observed across three other barnacle genomes (mean ENC ranging from 42.04 to 44.33; Table S6), highlighting a substantial reduction in the number of synonymous codon used in barnacles when compared to other metazoans (Figure 4A). A similar trend is observed when compared nucleotide composition in barnacle coding sequences when compared to other metazoans. All four barnacle genomes studied showed coding sequence GC content of *≈* 64%, 10 to 20% higher than the other studied genomes (Figure 4B; Table S6). The higher coding GC proportion in barnacles is also observed when looking at the GC proportion at 3rd codon positions and at 4-fold degenerate sites (Figure S6).

**Figure 4:**
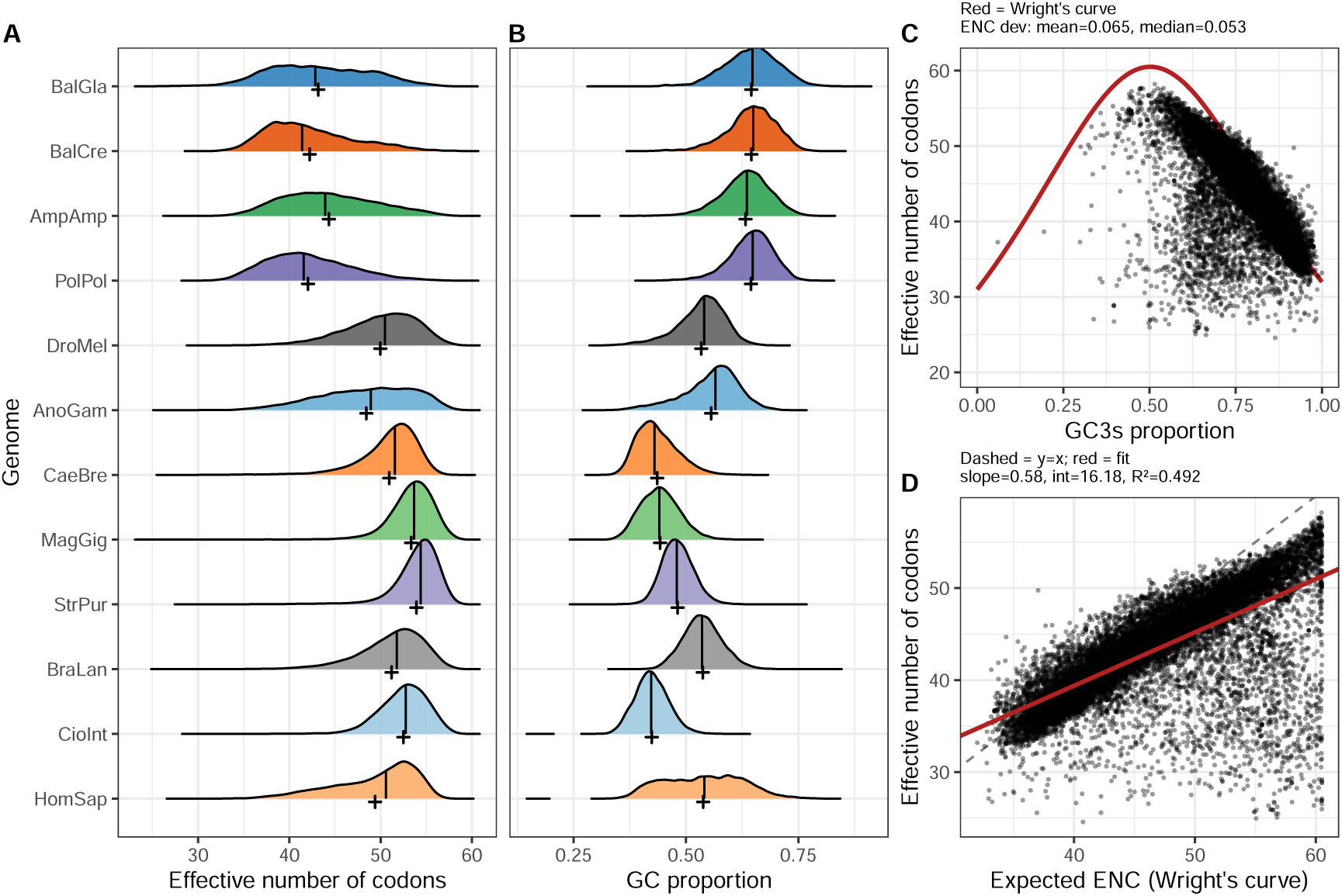
Codon usage biases in *Balanus glandula*. **A.** Genome-wide distribution of the effective number of codons (ENC) in *B. glandula* (BalGla) and other metazoan genomes (see Table S6), including the barnacles *B. crenatus* (BalCre), *A. amphitrite* (AmpAmp), and *P. pollicipes* (PolPol). The vertical lines show the 50^th^ quantile (median) of the distribution, while the crosses show the mean value. **B.** Distribution in the proportion of GC bases in coding sequences of 12 metazoan genomes. Vertical lines and crosses show the mean and median values, respectively. **C.** Relationship between the GC proportion at the 3rd codon site (GC3s) and ENC in *B. glandula*. Each black dot represents a gene, while the red curve shows the expected relationship in accordance with Wright’s curve (Wright, 1990). Points below the curve show genes with a smaller than expected number of effective codons, candidates for translational selection. **D.** Relationship between the expected number of effective codons (according to Wright’s curve) and the empirical ENC distribution in *B. glandula*. Red line shows the linear regression between expected and observed ENC distributions.

We explored the selective basis of the increased bias in codon usage in barnacle genomes by comparing the empirical ENC distribution against a theoretical expectation. In its original definition of the ENC statistic, Wright (1990) devised a relationship between GC3s and ENC, resulting in an expected number of effective codons for a given GC content. Deviations from this expectation, specifically observing a smaller number of effective codons than expected, suggest the presence of strong translational selection acting on the genome. After testing this relationship across these 12 genomes, we observe that the reduction in the number of effective codons observed in barnacles can in most cases be explained by their coding GC proportion (Figure S7; Figure S8), suggesting a codon usage bias driven by neutral processes. In *B. glandula* specifically, we observed a moderate deviation when comparing the empirical ENC distribution against Wright’s curve (Figure 4C); however, when comparing the linear relationship between observed and expected ENC, a better fit for GC biased genomes such as *B. glandula*, we do observe a reduction in the model’s slope and coefficient of determination (R^2^; Figure 4D). This poorer fit of the model indicates a smaller than expected number of effective codons, and suggests the presence of strong translational selection acting on the genome and the potential non-neutrality of at least some synonymous codons in large *B. glandula* populations.

Comparing the ENC and GC3s distribution in relation to population size highlights some interesting patterns. Evidence for selection-driven codon usage biases is observed for some hyper-diverse, large population species, e.g., the nematode *Caenorhabditis brenneri*, the Pacific oyster *Magallana gigas*, and the European lancelet *Branchiostoma lanceolatum* (Figure S7; Figure S8; Table S7). More moderate deviation is observed for other diverse species, e.g, the malaria mosquito *Anopheles gambiae* and the tunicate *Ciona intestinalis*. In contrast, in three of our studied barnacles (*B. crenatus*, *A. amphitrite*, and *P. pollicipes*), the observed codon usage biases appear are indistinguishable from neutral compositional processes. In *B. glandula*, evidence for non-neutral codon usage biases is moderate to strong depending on the method used. Overall, while we do not observe a clear relationship between population size and codon usage biases in the 12 studied genomes, the lineage-specific patterns described here provide an unexplored avenue of study of the efficacy and breadth of selection in highly diverse genomes.

### Barnacle populations exhibit elevated genetic polymorphism

We sequenced the whole genomes of eight *B. glandula* individuals from the central Oregon coast using Illumina short reads. After removal of two samples with poor quality sequences, and the stringent filtering of sites with low coverage, low quality, and/or missing genotypes, we recovered a total of 111.6 million genotyped sites (Table S4), 21.4 million (19.16%) of which were variant in the six retained samples. Among these variant sites, we recovered 20.5 million point mutations, 4.86% of which were multiallelic in just these six individuals. In total, the amount of polymorphism observed in our sampling translates to a mean density of 41.1 variant sites per kbp (median: 29, standard deviation: 40.82). Moreover, 1.1 million of these variant sites (5.23%) were located in coding sequences. These coding variants included 360 thousand missense (non-synonymous) mutations, corresponding to a mean of 18.93 non-synonymous variants per transcript (median: 10, standard deviation: 28.50). Overall, these represent a substantial proportion of coding variation segregating in central Oregon. This *B. glandula* population displayed a mean genome-wide nucleotide diversity (*π*) of 0.0506 (median: 0.0531, standard deviation: 0.0188; Figure 5; Table S8). For context, this magnitude of *π* is approximately 4*×* higher than that observed in *Anopheles gambiae* mosquitos, a species known for their high levels of polymorphism, *≈* 7*×* higher than the diversity in Sub-Saharan populations of *Drosophila melanogaster*, and *≈* 48*×* higher than the diversity observed in large human cohorts (Figure S9).

**Figure 5:**
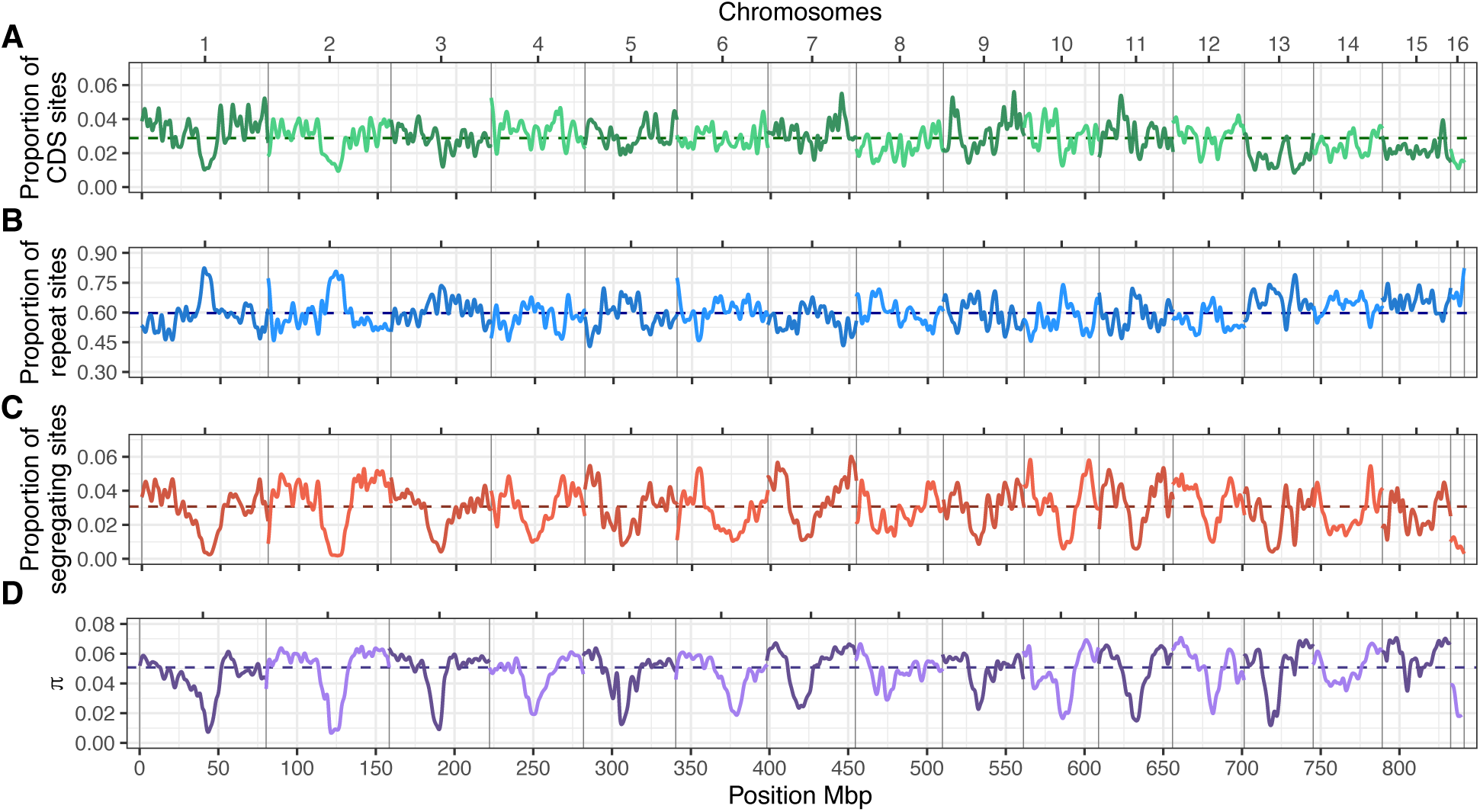
Nucleotide diversity (*π*) and coding and repeat density along the *Balanus glandula* genome. **A.** Density of coding sequences (CDS) along the genome, defined as the proportion of bases annotated as CDS within 10 kbp windows. Line reflects the kernel-smoothed values along a 1 Mbp window. Dashed horizontal line shows the genome-wide average proportion of CDS bases (0.0288). Top and bottom horizontal axes show the chromosome number and genomic position, respectively. **B.** Density of repeat elements annotated within 10 kbp windows, smoothed along 1 Mbp intervals. Dashed horizontal line shows the genome-wide average proportion of repeat bases (0.597). **C.** Density of segregating sites, calculated as the proportion of SNPs within 10 kbp windows, smoothed along 1 Mbp intervals. Dashed horizontal line shows the genome-wide average proportion of segregating SNPs per 10 kbp (0.0307). **D.** *π* within 10 kbp windows, smoothed along 1 Mbp intervals. Horizontal dashed line shows the mean genome-wide *π* (0.0506).

Along each chromosome, we did not observe any strong correlation between the density of specific genomic features (e.g., CDSs, lncRNAs, tRNAs, introns, and repeats) and *π*, particularly not at the scale of 10 kbp (Figure 5; Figure S10). However, stronger relationships are likely present at other genomic intervals given the differences in diversity seen across genetic elements (see below). The strongest correlation was, expectedly, observed between the density of segregating sites within 10 kbp windows and *π* (Spearman’s *ρ*: 0.706; Figure S10). The proportion of segregating sites in itself was not correlated with the density of particular genomic features, except for repeats; however, this is likely an artifact of the masking of genotypes located within repetitive intervals in the reference (see Population genetics of central Oregon barnacles). Despite the lack of clear correlation, the patterns of *π* observed along the chromosomes are qualitatively similar to those expected under regimes of strong linked selection, in which regions of reduced diversity are the result of differences in the recombination landscape (e.g., reduction in recombination in centromeric regions). Describing recombination in detail this species should thus be a focus of future studies in order to characterize how highly efficacious selection and linkage shape genetic diversity along the genome in *B. glandula*.

Looking at diversity across the genome, we observed an overall reduction in *π* at putatively functional sites. For example, the mean *π* in sequences spanning annotated CDSs is 0.0302 (median: 0.0270, standard deviation: 0.0221; Figure 6; Table S8), which represents a *≈* 1.6*×* reduction in diversity when compared to the genome-wide average. A reduction in diversity of similar magnitude is also observed in the sequences of lncRNAs (1.4*×*) and 5’ UTRs (1.3*×*). The largest reduction in diversity when compared to the genome-wide background was observed in tRNAs, with *π ≈* 4.3*×* lower than the genomic background and consistent with the strong functional constraint evidenced by their enrichment in highly conserved elements. In contrast to most other elements of the genome, introns display a relative increase in diversity, with a mean *π* of 0.0646 (median: 0.0650, standard deviation: 0.0299), approximately 1.3*×* higher than the genome-wide average.

**Figure 6:**
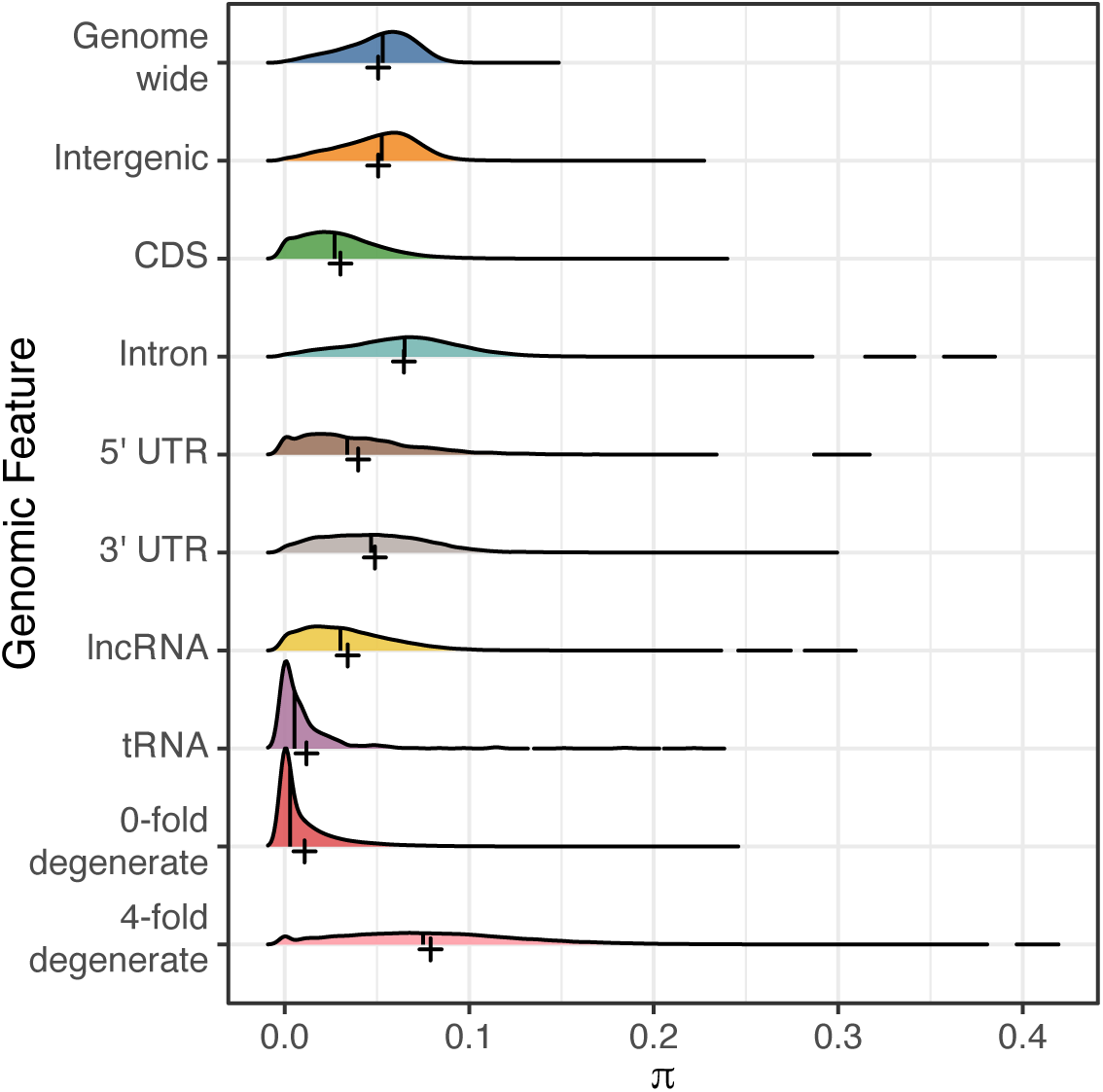
Nucleotide diversity (*π*) across genomic features in Oregon *Balanus glandula*. The density histogram shows the distribution of windowed *π* values across different genomic features annotated in the *B. glandula* assembly. The vertical lines show the 50^th^ quantile (median) of the distribution, while the crosses show the mean value.

Zooming in further on coding sequences, we explored the patterns of *π* at 0-fold degenerate (*π*_0_), in which any mutation produces a non-synonymous substitution, and 4-fold degenerate sites (*π*_4_), in which any mutation results in a synonymous change. This comparison allows us to compare diversity at constrained versus neutral (or putatively neutral) sites. In this central Oregon population of *B. glandula*, the mean *π*_0_, was 0.0107 (median: 0.00292, standard deviation: 0.0178). This is the lowest diversity among the studied categories in the genome, not a surprising result given the higher functional constraint expected at these sites. In contrast, mean *π*_4_ was 0.0790 (median: 0.0750, standard deviation: 0.0483), *≈* 1.6*×* higher than the genome-wide average and the highest in the genome. The nearly 8-fold difference between *π*_0_ and *π*_4_ highlights the strong constraint on non-synonymous sites and suggests that purifying selection is highly effective in this population, consistent with theoretical predictions for species with large effective population sizes.

### *Balanus glandula* exhibits large effective population sizes

We estimated the effective population size (N_e_) for a central Oregon *B. glandula* population sample using the whole-genome genotypes. As a simple estimate of the long-term N_e_ of this population, we first calculated N_e_ using Watterson’s estimator of genetic diversity (*θ*_W_), observing a mean genome-wide *θ*_W_ of 0.0617 (median: 0.0635, standard deviation: 0.0224; Figure S11). Assuming a per-base, per-generation mutation rate (*µ*) of 3 *×* 10*^−^*^9^ (as estimated by Nunez et al., 2020 for the northern acorn barnacle *S. balanoides*) and using the relationship between N_e_ and *θ*_W_, this yields a mean effective size of 5,137,172 (median: 5,289,211, standard deviation: 1,862,683), which is consistent with the expectation of large N_e_ in this species.

Additionally, we estimated N_e_ through time using a coalescent-based approach with msmc2 (Schiffels & Wang, 2020). This estimate also placed the historical effective size of the population on the order of 10^6^ individuals (between 10^6^ and 10^7^ generations ago) (Figure S12); however, given our small sample size and the use of only within sample pairwise comparisons, we were unable to obtain accurate N_e_ estimates at more recent (*<* 10^5^ generations) timescales using msmc2. Nevertheless, even these conservative estimates place *B. glandula* among the largest known effective population sizes for any eukaryote, yet orders of magnitude below the census sizes observed in the field — a disparity central to Lewontin’s Paradox and one we return to in the Discussion.

## Discussion

### Genome evolution in large populations

Evolutionary processes at large demographic scales are expected to leave distinct signatures in the genome, several of which are observed in our Pacific acorn barnacle assembly. For example, large populations are expected to harbor increased levels of segregating polymorphism, the product of both the reduction in the stochastic loss due to genetic drift and the increase in the population-scaled mutation rate. In accordance, we observe a barnacle genome with extreme levels of genetic diversity, both at the individual and population level (Figure S1; Table S4), a substantial proportion of which is located in coding sequences.

Similarly, large populations are expected to experience an increase in the efficacy of purifying selection, which is expected to reduce the levels of diversity and increase conservation across functional sequences. We see these patterns reflected in the *B. glandula* genome. Putatively functional sites, particularly coding exons and tRNAs, show both increased levels of sequence conservation (Figure 2) and a reduction in population-level genetic diversity (Figure 6; Table S8) compared to less constrained features of the genome. Notably, the rank ordering of the enrichment of conserved elements, with tRNAs and CDSs as the most enriched, parallels findings in insects (Siepel et al., 2005), suggesting that the targets of purifying selection are broadly conserved across arthropods despite the much larger and more repeat-rich genome of *B. glandula*. Increased efficacy of purifying selection is also reflected in the results of the McDonald-Kreitman test (Figure 3), where we observe a relative decrease in the proportion of non-synonymous polymorphism in a central Oregon *B. glandula* population, suggesting that weakly deleterious variants are more efficiently purged by selection (Y. F. Li et al., 2008). Positive selection is also expected to increase in efficacy in large populations, resulting in the fixation of a larger fraction of beneficial non-synonymous mutations. We observe this pattern as well: in this central Oregon population, the majority (54 to 61%, depending on the metric of *α* used) of non-synonymous substitutions reached fixation due to positive selection, a proportion higher than that observed in diverse *Drosophila* populations (Haller & Messer, 2017).

In large enough populations, selection should be efficacious enough that its effects extend to synonymous sites, where substitutions are normally considered neutral. This could result in a bias in the usage of synonymous codons, as different codons are favored over others due to GC biases, mRNA stability, tRNA abundance, and other factors (Galtier et al., 2018; Presnyak et al., 2015). These effects are observed in the *B. glandula* assembly in the form of reduction in the effective number of codons seen across protein-coding genes. On average, *B. glandula* uses *≈* 43 out of 61 available codons, a greater reduction than that observed in other metazoan genomes (Figure 4; Table S6). There is moderate evidence that this bias is not the product of compositional neutral processes (Figure S7; Figure S8), suggesting an increase in the efficacy of natural selection due to the large size of Pacific acorn barnacle populations. Moreover, these biases put into question the neutrality of at least some synonymous substitutions in this system, which could have important ramifications for the study of molecular evolution; however, the extent of these effects remain presently unknown and should be the subject of future study.

In contrast, the distribution of transposable and repeat elements in the *B. glandula* assembly was unexpected, as we find a highly repetitive genome composed of over 60% repeats (Figure 5; Table S3). Previous work has shown that purifying selection plays a key role in the accumulation and fixation of transposable elements (Bourque et al., 2018; Langmüller et al., 2023; Mérel et al., 2025), following the expectation that repetitive sequences are, on average, slightly deleterious. Given the increased efficacy of selection we observed in the rest of the genome, our expectation was to find a relatively small proportion of repeats. However, the high repeat content observed in the genome does not necessarily reflect the low efficacy of selection in these populations. Some could be low-frequency, slightly deleterious variants in the process of being purged from the population. Other repeats might represent beneficial genetic variation, as transposable elements are known to play roles in adaptive evolution (Schrader & Schmitz, 2019). For example, repeat elements can act as regulators of gene expression (Rech et al., 2022; Trizzino et al., 2017), and as such may provide the regulatory plasticity required for survival in rapidly changing intertidal environments. Regardless of their role, disentangling the population-wide repeat landscape and its fitness effects is likely impossible with a single haploid reference assembly. Thus, this observation drives the need for larger comparative genomic studies across barnacle populations before we can fully understand the dynamics of genomic repeats and transposable elements in this and other large populations.

### Molecular evolution in large populations: reconciling d_N_/d_S_ and the McDonald-Kreitman test

One of the more striking patterns in our results is the apparent discordance between our two measures of selection on protein-coding genes. The average pairwise d_N_/d_S_ between *B. glandula* and *B. crenatus* is 0.206, indicating that the majority of coding sites are under purifying selection. Yet the McDonald-Kreitman test reveals a seemingly opposite pattern: *≈* 61% of non-synonymous substitutions fixed between these species were driven by positive selection (*α*_asym_ = 0.610). At first glance, these results appear contradictory: how can coding sequences be under strong purifying selection while simultaneously showing pervasive adaptive evolution?

The resolution lies in the fact that d_N_/d_S_ and the McDonald-Kreitman test measure fundamentally different quantities. The d_N_/d_S_ ratio averages across all non-synonymous sites in a gene, the vast majority of which are under functional constraint, resulting in a ratio well below one. In contrast, the McDonald-Kreitman test partitions changes into polymorphism and divergence, using synonymous changes as a neutral baseline to ask whether there is an *excess* of non-synonymous fixations relative to the neutral expectation. This test is therefore sensitive to the subset of non-synonymous mutations that escape purifying selection and reach fixation, even when most non-synonymous sites in a gene are constrained. Moreover, pairwise d_N_/d_S_ is calculated from a single sequence per species and therefore incorporates segregating polymorphisms into its divergence estimate. In species with many weakly deleterious non-synonymous variants segregating at appreciable frequencies, this can inflate the non-synonymous divergence and distort estimates of selection (Y. F. Li et al., 2008). By explicitly separating polymorphism from fixation, the McDonald-Kreitman test avoids this conflation.

Y. F. Li et al. (2008) compared the performance of d_N_/d_S_ and the McDonald-Kreitman test across human, *Drosophila*, and yeast genomes and found that the two approaches identify distinct but correlated sets of genes as targets of selection. Notably, the McDonald-Kreitman test was more powerful than d_N_/d_S_ only in *Drosophila*, the species with the highest polymorphism, where it identified nearly three times as many genes under positive selection. In the lower-diversity human and yeast datasets, the pattern was reversed. This finding suggests that the statistical power of the McDonald-Kreitman test is greatest in high-polymorphism species, making it particularly well suited for systems like barnacles where segregating variation is abundant.

The pattern we observe in *B. glandula* is thus consistent with theoretical expectations for large populations: strong purifying selection efficiently removes the majority of deleterious non-synonymous mutations, keeping d_N_/d_S_ low, while at the same time the increased efficacy of positive selection drives a disproportionate fraction of the remaining non-synonymous substitutions to fixation, producing a high *α*. This coupling of strong purifying and positive selection is a hallmark of evolution in large populations and has been observed in *Drosophila*, where *α*_asym_ is approximately 0.57 (Messer & Petrov, 2013). Our estimate of *α*_asym_ of 0.610 in *B. glandula* is comparable to or slightly higher than that in *Drosophila*, consistent with the expectation that the even larger effective population size in barnacles further increases the efficacy of both modes of selection.

### Barnacles as models for evolution at biological extremes

Our work shows that a Pacific acorn barnacle population from central Oregon exhibits genetic diversity of over 5% (mean genome-wide *π* of 0.0506). These levels of genetic diversity are higher than those observed for other model systems known for their relatively large population sizes. For example, the genome-wide ϖ we observe in this population is approximately seven-fold higher than that observed in sub-Saharan populations of *D. melanogaster* (Figure S9) (Langley et al., 2012; W. Li et al., 2025; Nunez et al., 2025; Pool et al., 2012). Indeed, even the *π* observed at 0-fold degenerate sites, the least diverse and among the most constrained sites in the genome, exceeds the genome-wide diversity observed in these *Drosophila* populations. While elevated, this level of diversity is not unique to barnacles. Similar magnitudes of genome-wide *π* (*≈* 10*^−^*^2^) have been observed in other metazoan taxa, including corals (Meziere et al., 2025), bivalves (S. Liu et al., 2025), mosquitoes (Kent et al., 2025; The *Anopheles gambiae* 1000 Genomes Consortium, 2017), nematodes (Teterina et al., 2023), and lancelets (Brasó-Vives et al., 2026). Moreover, as with other marine invertebrates, barnacle life history traits are likely direct drivers of the elevated polymorphism observed, as factors such as large census sizes, small propagule size, large brood size, long larval dispersal, and broad geographic distribution have been positively correlated with high genetic diversity in these organisms (Leffler et al., 2012; Romiguier et al., 2014).

The magnitude of diversity observed in *B. glandula*solidly places them among hyperpolymorphic populations, those exhibiting diversity above 5% (Cutter et al., 2013). This adds barnacles to a growing list of emerging hyperpolymorphic model systems, which are key to studying the limits of biological variation and molecular evolution in natural populations. Moreover, our observations of extreme genetic diversity also have important implications for the study of species boundaries. For example, recent work by R. Hart et al. (2025) suggests that levels of polymorphism above 1% neutral diversity in eukaryotes likely serve as biological limits for the definition of species. Barnacles and other hyperpolymorphic populations put these putative cutoffs into question. We observe levels of genome-wide diversity of over 5% in just a very small portion of the biological range of the Pacific acorn barnacle. Nucleotide diversity is even higher (*≈* 8%) at 4-fold degenerate, and putatively neutral, sites (Figure 6; Table S8). Given the species’ broader geographical and ecological distribution, genetic diversity is likely to increase when looking across a larger spatial sampling. This observation reiterates the importance of future studies of genetic diversity of this species over its whole geographic range, providing some insights on the mechanisms that maintain population and species boundaries in the presence of extreme genetic diversity.

Beyond species boundaries, the extreme diversity observed in barnacles also speaks to a long-standing question in population genetics: the relationship between effective and census population sizes. From our results on diversity, we observe that this barnacle population has an effective size of approximately 10^6^. However, these populations also show densities of 10^4^ individuals per square meter (Menge, 2000). Based on this estimate, the census size of the Pacific acorn barnacle population in central Oregon should be, conservatively, in the billions of individuals. While the general disparity between the effective and census size estimates has been highlighted for decades, i.e., Lewontin’s Paradox (Lewontin, 1974), the mechanisms underlying the differences have been long debated (J. H. Gillespie, 1999, 2000; Kimura, 1983; J. M. Smith & Haigh, 1974) and are still the subjects of study (Buffalo, 2021). Our work on the Pacific acorn barnacle provides a unique opportunity to empirically study this disparity and the biological processes that maintain it.

### Conclusion

In this work, we take one of the first looks at the genomic signatures of evolution in large populations using the Pacific acorn barnacle, *Balanus glandula*. In accordance with population genetics theory, we find evidence for the increased efficacy of selection in this large barnacle population, manifested as both purifying selection reducing diversity and maintaining conservation across functional sites, and pervasive positive selection driving adaptive divergence in protein-coding genes. Surprisingly, however, we find a highly repetitive genome, in apparent contrast to the expectation that purifying selection should limit the proliferation of repetitive sequences. At the population scale, we observe average genome-wide nucleotide diversity exceeding 5%, placing this species among hyperpolymorphic organisms. Yet, while this diversity implies large effective population sizes (*≈* 10^6^ individuals), it still pales in comparison to the estimated census sizes of this population, underscoring the longstanding disparity between effective and census population sizes. Together, these findings establish *B. glandula* as a promising emerging model for studying the mechanisms of evolution at biological extremes.

## Data availability

The raw sequencing data for the assembly and annotation of the *B. glandula* and *B. crenatus* genomes is available on NCBI under BioProject PRJNA1378260. Raw whole-genome sequencing data for central Oregon *B. glandula* samples is available on NCBI BioProject PRJNA1434535. Copies of the bioinformatic scripts used for analysis and the raw files used to generate this document are available on GitHub (https://github.com/kr-colab/balanus_genome_ms). Additional copies of the final assembly and annotation files for *B. glandula* and *B. crenatus* available on Dryad (https://doi.org/10.5061/dryad.pvmcvdp21).

## Acknowledgments

We thank Tina Arredondo and the University of Oregon Genomics & Cell Characterization Core Facility for their assistance with DNA extraction, library preparation, and sequencing. Also, we thank George von Dassow and Richard B. Emlet for their help with sample collection and species identification, Nathaniel S. Pope for assistance with the Ag1000G and 1kGP datasets, and Kira M. Long for her comments and feedback on the manuscript. Lastly, we thank members and collaborators of the Kern-Ralph co-lab for their assistance with sample collection, project feedback, and manuscript review. A.D.K. was supported in part by NIH grant R35GM148253.

## Author Contributions

A.G.R.-C. designed the experiments, collected samples, analyzed the data, and wrote the first draft of the manuscript. S.T.S. designed the experiments, collected samples, analyzed the data, and edited the manuscript. E.J. collected samples and edited the manuscript. J.P.W. edited the manuscript. A.D.K. oversaw the project, designed the experiments, collected samples, and analyzed the data. All authors read and approved the final version of the text.

