## Supplemental files for "Large population sizes shape genome evolution in barnacles"

August 17, 2026

#### Supplemental Methods

##### Assembly and annotation of *Balanus crenatus*

###### High-molecular weight DNA and PacBio sequencing

For genome sequencing using PacBio Hi-Fi reads, we sampled several *Balanus crenatus* individuals from the docks in the Oregon Institute of Marine Biology, in Charleston, Oregon, USA (43.346 N, 124.325 W) in November 2023 during low tide. High-molecular weight (HMW) DNA was extracted from the dissected testes using the PacBio Nanobind tissue kit (PacBio). DNA was assessed for quality by fluorometric quantification of concentration (Qubit; Invitrogen) and fragment length distribution (Fragment Analyzer; Advanced Analytical). The extracted DNA of a single individual was sequenced on two SMRTcells of a PacBio Sequel II machine at the University of Oregon's Genomics and Cell Characterization Core Facility (GC3F). These two sequencing runs yielded a total of 3.06 million reads with a read N50 of 14.8 kbp.

###### Estimating genome size and heterozygosity

We used $k$-mers to estimate the size and heterozygosity of the *B. crenatus* genome. First, we counted $k$-mers present in the PacBio HiFi reads using JELLYFISH version `2.2.10` (Marçais & Kingsford, 2011), counting 21-mers (`--mer-len 21`) present in both strands (`--canonical`). These counts were then used to generate an empirical distribution of 21-mers using the JELLYFISH `histo` command. The final model of the diploid, genome-wide $k$-mer distribution was then performed using GenomeScope2 version `2.0` (Ranallo-Benavidez et al., 2020). We estimated the *B. crenatus* genome size to be 755,696,144 bp in length, with a heterozygosity of 2.25%.

###### Contig-level genome assembly

We generated a contig-level genome assembly using HIFIASM version `0.19.8-r603` (Cheng et al., 2021, 2022), using strict parameters for the identification and purging of haplotig sequences. This assembly was generated using an estimated haploid genome size of 800 Mbp (`--hg-size 800m`), dropping $k$-mers observed over ten times the average coverage (`-D 10.0`), and allowing scaffolding (`--dual-scaf`). For handling haplotigs, we set the expected coverage at homozygous sites to 40x (`--hom-cov 40`), the upper bound of coverage to purging duplicates to 40x (`--purge-max 40`), and the similarity threshold between duplicate haplotigs to 25% (`-s 0.25`). We assessed the contiguity of this assembly using QUAST version `5.2.0` (Gurevich et al., 2013), showing an assembly composed of 2,932 contigs, with a total length of 1.20 Gbp, and a contig N50 of 746 kbp. Additionally, COMPLEASM version `0.12-r237` (Huang & Li, 2023) revealed this assembly 94.96% complete (with 19.64% duplicates) against the `arthropoda_odb10` reference ortholog dataset from BUSCO (Manni et al., 2021; Simão et al., 2015; Tegenfeldt et al., 2025).

##### Identifying contamination

We screened the *B. crenatus* contig-level assembly for contamination using MMseqs2 release `15-6f452` (Steinegger & Söding, 2017). After indexing the assembly, taxonomic assignment was done for each contig. As described in the MMseqs2 documentation, protein fragments were extracted across the six possible frames, which were then mapped against the NCBI `nr` and `SwissProt` databases. We retained only sequences assigned to the order Thecostraca (NCBI taxid `116172`). This process removed 236 fragments spanning $\approx$ 19.7 Mbp, producing an assembly with 2,696 contigs, a total size of 1.18 Gbp, and a contig N50 of 758.4 kbp. The resulting gene-completeness against the `arthropoda_odb10` reference set was 94.57% with 19.45% duplicates.

##### Purging haplotigs

After screening the assembly for contamination, we identified and purged haplotig sequences using PURGE_DUPS version `1.2.5` (Guan et al., 2020). We first mapped the PacBio HiFi reads to the contig-level assembly with MINIMAP2 `-x map-hifi`. From these alignments, we calculated the read-depth histogram using `pbcstat` and determined the base-level coverage cutoffs using `calcuts`. After splitting the assembly at gaps with `split_fa`, we performed an assembly self-alignment using MINIMAP2 `-x asm20`. Haplotigs were then marked according to the empirical coverage cutoffs using the `purge_dups` command. Lastly, the assembly FASTA was processed to remove sequences tagged as haplotigs using `get_seqs`, only removing sequences at the ends of contigs (`-e`) and allowing the splitting of contigs (`-s`). The purging reduced the number of contigs in the assembly to 1,548, resulting in a total assembly size of 907.4 Mbp, a contig N50 of 925.5 kbp, and a gene-completeness of 93.39% with 3.16% duplicates (against the `arthropoda_odb10` reference set).

##### Polishing and correcting

The purged contig-level assembly was then polished and self-corrected using INSPECTOR software version `1.0.1` (Y. Chen et al., 2021). First, we evaluated the assembly using `inspector.py`, mapping the PacBio HiFi reads against the assembly. The assembly was then corrected using `inspector-correct.py`. After polishing, the assembly contained 1,548 contigs, a total length of 907.3 Mbp, and a contig N50 of 925.6 kbp. Gene-completeness against the `arthropoda_odb10` reference set was 93.39% with 3.26% duplicates.

##### Reference-based scaffolding

We used a CACTUS whole-genome alignment (Armstrong et al., 2020) to scaffold the assembly. Using CACTUS version `2.9.7` we aligned the chromosome-level *B. glandula* assembly against the

*B. crenatus* contigs, exporting the output alignments in MAF format. After alignment, we used the RAGOUT version `2.3` software (Kolmogorov et al., 2018) to perform reference-based scaffolding, disabling the breaking of input sequences (`--solid-scaffolds`) and enabling the repeat resolution algorithm (`--repeats`). After running RAGOUT, we only kept the *B. crenatus* sequences that were successfully aligned to *B. glandula*. This scaffolded assembly was composed of 220 sequence fragments, with a total length of 851 Mbp, and a scaffold N50 of 5.58 Mbp. Additionally, the assembly was 89.64% gene-complete (with 2.37% duplicates) when compared to the `arthropoda_odb10` reference set.

##### Genome annotation

Repetitive elements were annotated in the scaffolded *B. crenatus* assembly using the EarlGrey software version `4.1.0` (Baril et al., 2024). When running EarlGrey, we provided the Crustacea Dfam 3.8 partition as an initial consensus library (`-l`), specified "arthropoda" as the search term for RepeatMasker version `4.1.2` (Smit et al., 2013), clustered the final TE library to reduce redundancy (`-c yes`), and generated a soft-masked FASTA as output (`-d yes`). Similar to *B. glandula*, the repeat annotation for *B. crenatus* revealed a highly repetitive genome, with 494.0 Mbp (58.03%) of the assembly masked. Moreover, the largest proportion of identified repeats remained unclassified, 289.4 Mbp or 40.0% of the assembly, further highlighting the lack of barnacle-representative sequences in public repeat databases.

Following the annotation of repeats, we annotated protein-coding sequences in the *B. crenatus* assembly by lifting over gene-models from the curated *B. glandula* annotation using LiftOn version `1.0.5` (Chao et al., 2024). LiftOn lifts over annotations between genomes by integrating both protein-to-genome alignments with MINIPROT (Li, 2023) and nucleotide-to-nucleotide alignments using Liftoff (Shumate & Salzberg, 2021). Given the high diversity observed within these genomes and the high divergence between them, we relaxed several of the LiftOn parameters to maximize the DNA and protein sequence identity scores, as well as maintaining a high BUSCO completeness. We included 10% of the flanking sequence of each gene as part of the alignment (`-f flank 0.1`), we allowed the software to identify gene copies with sequence similarity of at least 50% (`-copies -sc 0.5`), used a 25% coverage threshold for the mapping of parent (`-a 0.25`) and child sequences (`-s 0.25`), and enabled a post-liftover refinement of the transferred gene models (`-polish`). We also used the "general" splice model for MINIPROT (`-j 1`). This liftover successfully annotated 15,841 genes, including 15,904 protein-coding transcripts, in the *B. crenatus* assembly. While this is $\approx$ 5,000 fewer protein-coding genes than those annotated for *B. glandula*, this annotation was still 82.63% gene-complete, with 9.67% duplicates, when compared to the `arthropoda_odb10` reference ortholog set using COMPLEASM `protein`.

#### Processing annotations of published barnacle assemblies

##### ***Pollicipes pollicipes***

We used the published RefSeq annotation for the gooseneck barnacle *P. pollicipes* (NCBI accession: `GCF_011947565.3`; Bernot et al., 2022; Table S9). This annotation was processed with AGAT version `1.4.3` (Dainat, 2022) to filter out non-coding annotations, extract the longest transcript as representative for the gene, and standardize gene and sequence IDs for downstream analysis. This RefSeq annotation contains 20,420 protein-coding genes and is 88.72% gene-complete, with 35.93% duplicates, when validated against the `arthropoda_odb10` reference ortholog dataset using COMPLEASM `protein`.

##### *Amphibalanus amphitrite*

We combined the genome and the annotation across two different assemblies for the striped barnacle, *A. amphitrite*. First, we used the chromosome-level genome assembly for the species generated by Han et al. (2024). While the assembled sequences for this genome are available (NCBI accession: `GCA_037642225.1`; Table S9), no annotation has been published for this assembly. Therefore, we then took the published RefSeq annotation (NCBI accession: `GCF_019059575.1`; Table S9), based on the scaffold-level genome assembly by J.-H. Kim et al. (2019). We annotated the chromosome-level assembly by lifting over the RefSeq annotation using the LiftOn software version `1.0.5` (Chao et al., 2024), allowing the software to identify gene copies with a sequence similarity of 80% (`-copies -sc 0.8`), and using the "general" splice model in miniprot (`-j 1`). This liftover process successfully annotated 23,035 protein-coding genes. The annotation was then processed with agat version `1.4.3` to extract the longest transcript as representative for the gene and to standardize gene and sequence IDs for downstream analysis. This processed annotation was then validated using compleasm `protein` using the `arthropoda_odb10` reference ortholog set, showing it was 88.08% complete with 12.24% duplicates.

##### *Amphibalanus improvisus*

We annotated the chromosome-level assembly of the Bay barnacle, *A. improvisus* (NCBI accession: `GCA_964274985.1`; Bishop et al., 2025; Table S9) using publicly available transcriptomic data. We downloaded two public RNAseq datasets for this species, from NCBI BioProjects `PRJNA528777` and `PRJNA528169` (Table S10; Abramova et al., 2019). The raw reads were then processed using fastp version `0.23.4` (S. Chen, 2023; S. Chen et al., 2018), and aligned to the *A. improvisus* assembly using hisat2 version `2.2.1` (D. Kim et al., 2019). We then used BRAKER version `3.0.8` (Gabriel et al., 2024) to annotate the genome using the evidence from both the RNAseq alignments and the Arthropod representative protein sequences from OrthoDBv11 (Kriventseva et al., 2019; Zdobnov et al., 2021). In BRAKER, we enforced the recovery of orthologs from the Arthropod BUSCO dataset in the annotation (`--busco_lineage=arthropoda_odb10`). After running BRAKER, we processed the resulting annotation using agat to extract the longest transcript as representative for the gene and to standardize gene and sequence IDs for downstream analysis. This process annotated 16,098 protein-coding genes, and was 96.05% gene-complete, with 16.36% duplicates, after validation with compleasm against the `arthropoda_odb10` reference ortholog dataset.

##### *Capitulum mitella*

We annotated the publicly-available chromosome-level genome assembly of the pedunculate barnacle, *C. mitella* (NCBI accession: `GCA_030062745.1`; D. Chen et al., 2021; Table S9) using RNAseq short read data publicly available on the NCBI SRA database (Table S10). We followed the same methodology used for *A. improvisus*: processing the short reads with fastp, aligning to the genome with hisat2, annotating with BRAKER, and processing the resulting annotation with agat. This approach resulted in the annotation of 10,504 protein-coding genes, which produced an annotation that was 97.53% gene-complete, with 14.02% duplicate sequences when compared to the `arthropoda_odb10` reference ortholog dataset. However, given the comparatively smaller number of annotated sequences, and the resulting reduction in the number of orthologous sequences recovered when incorporating this species, we excluded *C. mitella* from most downstream analyses.

#### Comparing barnacle genetic diversity against model systems

##### *Drosophila melanogaster*

We calculated nucleotide diversity ($\pi$) in sub-Saharan populations of the fruit fly *D. melanogaster* to establish a baseline for comparison against our central Oregon barnacle population. To achieve this, we first obtained the sequencing data for phase 2 of the *Drosophila* Population Genomics Project (DPGP2; Lack et al., 2016; Pool et al., 2012). We processed the aligned FASTA files for all sub-Saharan samples (the 'RG' population; N=27) to generate a VCF using the SNP-SITES version `2.5.1` software (Page et al., 2016). We exported both variant and invariant sites (`-b`) for downstream compatibility. The resulting gVCFs were then processed using a custom AWK command and BCFTOOLS version `1.21` (Danecek et al., 2021) to recode deletions, remove missing alleles, and remove sites with over 25% missing genotypes. Nucleotide diversity along 10 kbp windows was then calculated from the filtered VCF using PIXY version `2.0.0.beta12` (Bailey et al., 2025; Korunes & Samuk, 2021), allowing multiallelic sites to be used in the calculation (`--include_multiallelic_snps`). For comparing average $\pi$ across species, we removed any window with less than 10% of available sites.

##### *Anopheles gambiae*

Comparing against another highly diverse invertebrate, we obtained genotypes from phase 3 of the *Anopheles gambiae* 1000 Genomes Project (Ag1000G; The *Anopheles gambiae* 1000 Genomes Consortium, 2017, 2021). The genotypes in ZARR format were loaded as a per-chromosome `HaplotypeMatrix` in the software PG_GPU version `0.1.1` (Pope et al., 2026). Diversity was calculated on a subset of Ag1000G samples from Bangui in the Central African Republic (N=55). We applied an accessibility mask to remove both low-quality sites and to remove variants located in known *An. gambiae* inversions (e.g., 2La, 2Rb, 2Rc). We then used PG_GPU's windowed statistics (`windowed_analysis`) to calculate $\pi$ (`statistics=["pi"]`) in non-overlapping 10 kbp windows (`window_size=10000`, `step_size=10000`), accounting for the presence of masked sites (`span_normalize=True`). For comparing average $\pi$, windows with less than 10% of available sites were removed.

##### *Homo sapiens*

Similarly, we compared genetic diversity in *B. glandula* against a large human cohort. For this, we used the human haplotypes from the high-coverage whole-genome sequencing release of the 1000 Genomes Project (1kGP; Byrska-Bishop et al., 2022). Similar to the *An. gambiae* analysis, we first loaded the 1kGP data as a `HaplotypeMatrix` in PG_GPU. For the diversity calculation, we subsampled haplotypes to only includes Yoruba individuals (YRI), removing and removing any parent-offspring pairs according to the available pedigree information. In total, we retained 66 YRI individuals. We applied an accessibility mask to retain only high-mappability and non-duplicated regions of the genome, in accordance to the publicly available Genome-in-a-Bottle (GIAB) genome stratification features (GIAB version `3.6`; Dwarshuis et al., 2024). Then, we calculated $\pi$ in 10 kbp non-overlapping windows using PG_GPU's windowed analysis. For comparing average $\pi$ across species, windows with less than 10% of available sites were removed.

### Supplemental Tables

Table S1: **Assembly statistics for the *Balanus glandula* genome at different stages of the assembly process.** Default contigs refers to the sequences assembled by HIFIASM with default parameters. Optimized contigs denotes the sequences generated using HIFIASM followed by removal of contaminant sequences and purging of haplotigs. Hi-C scaffolds refer to the scaffolds generated by YAHS. Curated assembly refers to the sequences generated after manual curation of the Hi-C contact map and polishing with INSPECTOR. BUSCO results refer to the `arthropoda_odb10` dataset.

| Assembly Stage | Default contigs | Optimized contigs | Hi-C scaffolds | Curated assembly |
|---|---|---|---|---|
| Sequence length (Mbp) | 1.64 | 1.05 | 1.05 | 1.04 |
| Fragments (N) | 4,212 | 1,342 | 650 | 592 |
| Contig N50 (Mbp) | 0.734 | 1.35 | 1.18 | 1.18 |
| Scaffold N50 (Mbp) | `NA` | `NA` | 50.96 | 51.41 |
| BUSCO complete (%). | 95.66 | 93.88 | 93.98 | 94.08 |
| BUSCO single (%) | 45.31 | 89.04 | 89.24 | 89.44 |
| BUSCO duplicates (%) | 50.35 | 4.84 | 4.74 | 4.64 |
| BUSCO fragmented (%) | 0.79 | 1.18 | 0.89 | 0.89 |
| BUSCO missing (%) | 3.55 | 4.94 | 5.13 | 5.03 |
| BUSCO size (N) | 1,013 | 1,013 | 1,013 | 1,013 |

Table S2: **Completeness metrics for the *Balanus glandula* protein-coding gene annotation.** BUSCO completeness was determined by COMPLEASM `protein`.

| Metric | `arthropoda_odb10` | `crustacea_odb12` |
|---|---|---|
| BUSCO complete (%) | 93.58 | 84.77 |
| BUSCO single (%) | 79.76 | 72.66 |
| BUSCO duplicates (%) | 13.82 | 12.11 |
| BUSCO fragmented (%) | 1.78 | 4.10 |
| BUSCO missing (%) | 4.64 | 11.13 |
| BUSCO size (N) | 1,013 | 1,536 |

Table S3: **High-level repeat annotation of the *Balanus glandula* assembly.** Repeats were first identified with the EarlGrey software. The classification is the merge of EarlGrey's homology-based assignment, followed by the re-classification of unknown elements by TEsorter and DeepTE. Coverage is the total number of masked sites per class; "% Genome" is the coverage divided by assembly size (1,043,519,078 bp).

| Classification | Coverage (bp) | Count | % Genome | Distinct elements |
|---|---|---|---|---|
| DNA | 379,178,613 | 895,512 | 36.34 | 35 |
| LINE | 107,103,442 | 223,314 | 10.26 | 20 |
| LTR | 50,161,694 | 120,887 | 4.81 | 8 |
| Other (Simple Repeat, Microsatellite, RNA) | 37,359,678 | 74,786 | 3.58 | 3 |
| Penelope | 25,833,119 | 65,921 | 2.48 | 4 |
| Rolling Circle | 22,238,826 | 42,257 | 2.13 | 1 |
| Retroposon | 2,095,328 | 5,834 | 0.20 | 2 |
| SINE | 1,253,619 | 4,437 | 0.12 | 3 |
| DIRS | 1,184,091 | 1,285 | 0.11 | 2 |
| Unclassified | 61,450,725 | 148,647 | 5.89 | 1 |
| Total | 687,859,135 | 1,582,880 | 65.92 | 79 |

Table S4: **Genetic variation in central Oregon *Balanus glandula*.**

| Metric | Reference Individual | Cape Perpetua (N=6) |
|---|---|---|
| Reference sequence length (bp) | 904,940,360 | 904,940,360 |
| Genotyped sites post-filtering (bp) | 314,105,136 | 111,668,206 |
| Total variant sites (bp) | 9,081,889 | 21,395,217 |
| Percent variant sites (%) | 2.89 | 19.16 |
| SNPs (N) | 8,725,877 | 20,501,788 |
| Indels (N) | 356,012 | 893,429 |
| Multiallelic SNPs (N) | NA | 996,103 |
| Percent multiallelic SNPs (%) | NA | 4.86 |
| Mean variant sites per kbp (N) | 14.66 | 41.05 |
| Median variant sites per kbp (N) | 6 | 29 |
| St Dev variant sites per kbp (N) | 19.86 | 40.82 |
| Variant sites in CDSs (N) | 398,773 | 1,118,874 |
| Percent variants in CDS (%) | 4.69 | 5.23 |
| Mean variants per CDS (N) | 4.34 | 12.19 |
| Median variants per CDS (N) | 1 | 6 |
| St Dev variants per CDS (N) | 11.75 | 25.84 |

Table S5: **Enchrichment of annotated features among phastCons highly conserved elements.** The total feature proportion was determined according to the 841,166,098 bp corresponding to the 16 chromosome-level scaffolds. Proportion of highly-conserved elements (HCE) was calculated based on the 50,849,898 bp-span of all HCEs in the genome. The number of chromosomes denotes the number of chromosome-level scaffolds containing annotations of the given feature.

| Feature | Total feature length (bp) | Total feature proportion | HCE feature length (bp) | HCE feature proportion | Mean $\log_2$ enrichment | Median $\log_2$ enrichment | St Dev $\log_2$ enrichment | Number of chromosomes |
|---|---|---|---|---|---|---|---|---|
| 3’ UTR | 2,358,104 | 2.80e−3 | 152,598 | 3.00e−3 | −0.0861 | 0.0332 | 0.508 | 16 |
| 5’ UTR | 530,151 | 6.30e−4 | 27,679 | 5.44e−4 | −0.292 | −0.332 | 0.453 | 16 |
| CDS | 16,273,411 | 0.0193 | 2,225,138 | 0.0438 | 0.752 | 0.984 | 0.926 | 16 |
| TE | 370,573,528 | 0.441 | 17,505,920 | 0.344 | −0.359 | −0.305 | 0.180 | 16 |
| intergenic | 196,612,776 | 0.234 | 10,688,761 | 0.210 | −0.167 | −0.178 | 0.167 | 16 |
| intron | 111,562,654 | 0.133 | 8,575,960 | 0.169 | 0.419 | 0.297 | 0.432 | 16 |
| lncRNA | 2,211,027 | 2.63e−3 | 130,744 | 2.57e−3 | −0.174 | −0.133 | 0.567 | 16 |
| multiple | 141,022,121 | 0.168 | 11,537,936 | 0.227 | 0.418 | 0.438 | 0.167 | 16 |
| rRNA | 2,106 | 2.50e−6 | 0.000 | 0.000 | NaN | NaN | NaN | 6 |
| tRNA | 20,220 | 2.40e−5 | 5,162 | 1.02e−4 | 2.143 | 2.041 | 0.959 | 16 |

Table S6: **Codon usage biases across metazoan genomes.** Table shows the codon usage statistics (effective number of codons; ENC), GC proportion, and GC proportion at third codon positions (GC3s) for length-filtered (CDS $\geq$ 300 bp, within 2 standard deviations of the per-species mean length) genes in each genome. The first four rows (`BalGla`, `BalCre`, `AmpAmp`, and `PolPol`) denote the focal barnacle genomes.

| | | | ENC | | | GC | | | GC3s | | |
|---|---|---|---|---|---|---|---|---|---|---|---|
| Genome | Species | Number of Genes | Mean | Median | StDev | Mean | Median | StDev | Mean | Median | StDev |
| `BalGla` | *Balanus glandula* | 16,217 | 43.17 | 42.84 | 5.52 | 0.645 | 0.648 | 0.061 | 0.781 | 0.796 | 0.102 |
| `BalCre` | *Balanus crenatus* | 7,675 | 42.23 | 41.40 | 4.78 | 0.645 | 0.649 | 0.055 | 0.815 | 0.837 | 0.093 |
| `AmpAmp` | *Amphibalanus amphitrite* | 25,242 | 44.33 | 43.93 | 5.05 | 0.632 | 0.636 | 0.054 | 0.775 | 0.792 | 0.099 |
| `PolPol` | *Pollicipes pollicipes* | 17,607 | 42.04 | 41.57 | 4.56 | 0.644 | 0.648 | 0.049 | 0.829 | 0.845 | 0.082 |
| `DroMel` | *Drosophila melanogaster* | 12,312 | 49.97 | 50.47 | 3.99 | 0.534 | 0.540 | 0.053 | 0.630 | 0.640 | 0.110 |
| `AnoGam` | *Anopheles gambiae* | 11,804 | 48.43 | 48.92 | 5.09 | 0.556 | 0.566 | 0.062 | 0.667 | 0.688 | 0.136 |
| `CaeBre` | *Caenorhabditis brenneri* | 22,904 | 50.95 | 51.55 | 3.21 | 0.436 | 0.430 | 0.045 | 0.411 | 0.399 | 0.088 |
| `MagGig` | *Magallana gigas* | 24,963 | 53.34 | 53.66 | 2.58 | 0.442 | 0.441 | 0.043 | 0.433 | 0.426 | 0.085 |
| `StrPur` | *Strongylocentrotus purpuratus* | 25,759 | 53.92 | 54.39 | 2.68 | 0.481 | 0.480 | 0.035 | 0.489 | 0.486 | 0.079 |
| `BraLan` | *Branchiostoma lanceolatum* | 23,301 | 51.20 | 51.78 | 3.57 | 0.537 | 0.536 | 0.043 | 0.620 | 0.619 | 0.095 |
| `CioInt` | *Ciona intestinalis* | 14,370 | 52.49 | 52.75 | 2.66 | 0.424 | 0.422 | 0.037 | 0.369 | 0.367 | 0.069 |
| `HomSap` | *Homo sapiens* | 21,573 | 49.39 | 50.57 | 4.51 | 0.538 | 0.541 | 0.087 | 0.593 | 0.603 | 0.163 |

Table S7: **Summary of ENC model fits across metazoan genomes.** Table shows the fits for the model between the expected and observed effective number of codons (ENC) across 12 metazoan genomes. ENC deviation values represents the deviation from Wright's curve, as shown in Figure S7. Larger ($\gg 0$) values denote a reduction in the observed number of effective codons when compared to the GC3s-based expectation, suggesting translational selection. The linear regression values represent the linear relationship between the observed and expected number of effective codons, as seen in Figure S8. Deviations in ENC product of translational selection are seen as a reduction in both slope and coefficient of determination ($R^2$).

| | | | ENC deviation | | | | | Linear regression | | |
|---|---|---|---|---|---|---|---|---|---|---|
| Genome | Species | Number of Genes | Mean | Median | StDev | Min | Max | Slope | Intercept | $R^2$ |
| `BalGla` | *B. glandula* | 16,217 | 0.065 | 0.053 | 0.090 | −0.344 | 0.571 | 0.580 | 16.18 | 0.492 |
| `BalCre` | *B. crenatus* | 7,675 | 0.040 | 0.044 | 0.052 | −0.235 | 0.382 | 0.725 | 10.19 | 0.844 |
| `AmpAmp` | *A. amphitrite* | 25,242 | 0.052 | 0.055 | 0.047 | −0.225 | 0.517 | 0.747 | 9.27 | 0.868 |
| `PolPol` | *P. pollicipes* | 17,607 | 0.025 | 0.032 | 0.054 | −0.265 | 0.436 | 0.746 | 9.77 | 0.806 |
| `DroMel` | *D. melanogaster* | 12,312 | 0.092 | 0.092 | 0.039 | −0.125 | 0.356 | 0.754 | 8.44 | 0.760 |
| `AnoGam` | *A. gambiae* | 11,804 | 0.074 | 0.075 | 0.040 | −0.184 | 0.480 | 0.779 | 7.58 | 0.871 |
| `CaeBre` | *C. brenneri* | 22,904 | 0.107 | 0.100 | 0.057 | −0.079 | 0.518 | 0.436 | 26.00 | 0.175 |
| `MagGig` | *M. gigas* | 24,963 | 0.079 | 0.076 | 0.040 | −0.108 | 0.567 | 0.560 | 20.89 | 0.330 |
| `StrPur` | *S. purpuratus* | 25,759 | 0.088 | 0.083 | 0.037 | −0.287 | 0.465 | 0.853 | 3.51 | 0.358 |
| `BraLan` | *B. lanceolatum* | 23,301 | 0.085 | 0.082 | 0.046 | −0.166 | 0.487 | 0.642 | 15.17 | 0.624 |
| `CioInt` | *C. intestinalis* | 14,370 | 0.060 | 0.057 | 0.038 | −0.156 | 0.428 | 0.591 | 19.45 | 0.588 |
| `HomSap` | *H. sapiens* | 21,573 | 0.086 | 0.091 | 0.046 | −0.201 | 0.486 | 0.693 | 11.83 | 0.818 |

Table S8: **Nucleotide diversity ($\pi$) across different genomic features.**

| Feature | $\pi$ Mean | $\pi$ Median | $\pi$ StDev | Number of windows | Window length (bp) Mean | Window length (bp) Median | Window length (bp) StDev | Number of sites | Sites per window Mean | Sites per window Median | Sites per window StDev |
|---|---|---|---|---|---|---|---|---|---|---|---|
| Genome wide | 0.0506 | 0.0531 | 0.0188 | 44,599 | 9,998.570 | 10,000 | 103.774 | 93,583,599 | 2,098.334 | 1,932 | 821.464 |
| Intergenic | 0.0507 | 0.0526 | 0.0210 | 29,219 | 7,688.054 | 10,000 | 3,531.991 | 45,677,153 | 1,563.269 | 1,431 | 956.166 |
| CDS | 0.0302 | 0.0270 | 0.0221 | 78,630 | 270.688 | 161 | 405.314 | 9,447,233 | 120.148 | 80 | 163.758 |
| Intron | 0.0646 | 0.0650 | 0.0299 | 67,638 | 2,182.716 | 686 | 5,257.832 | 34,345,262 | 507.781 | 209 | 1,090.025 |
| 5' UTR | 0.0398 | 0.0338 | 0.0309 | 6,013 | 153.063 | 114 | 148.787 | 424,412 | 70.582 | 54 | 63.678 |
| 3' UTR | 0.0489 | 0.0468 | 0.0294 | 8,218 | 648.360 | 542 | 485.406 | 2,083,933 | 253.582 | 210 | 194.076 |
| lncRNA | 0.0341 | 0.0302 | 0.0246 | 27,168 | 399.958 | 184 | 511.811 | 4,557,588 | 167.756 | 91 | 200.540 |
| tRNA | 0.0118 | 5.21e−3 | 0.0236 | 430 | 66.177 | 72 | 16.982 | 13,804 | 32.102 | 32 | 14.305 |
| 0-fold degenerate | 0.0107 | 2.92e−3 | 0.0178 | 75,895 | 269.321 | 162 | 401.035 | 5,397,246 | 71.115 | 46 | 96.221 |
| 4-fold degenerate | 0.0790 | 0.0750 | 0.0483 | 39,170 | 281.364 | 176 | 380.879 | 1,410,361 | 36.006 | 23 | 46.585 |

Table S9: **Publicly available NCBI assemblies used for comparative analyses.** For the NCBI accessions, `GCF_*` denotes RefSeq assemblies and annotations. `GCA_*` accessions denote GenBank assemblies and (when available) annotations.

| Genome | Species | NCBI accession |
|---|---|---|
| AmpAmp | *Amphibalanus amphitrite* | GCF_019059575.1 |
| AmpAmp | *Amphibalanus amphitrite* | GCA_037642225.1 |
| AmpImp | *Amphibalanus improvisus* | GCA_964274985.1 |
| CapMit | *Capitulum mitella* | GCA_030062745.1 |
| PolPol | *Pollicipes pollicipes* | GCF_011947565.3 |
| DroMel | *Drosophila melanogaster* | GCF_000001215.4 |
| AnoGam | *Anopheles gambiae* | GCF_943734735.2 |
| CaeBre | *Caenorhabditis brenneri* | GCA_964036135.1 |
| MagGig | *Magallana gigas* | GCF_963853765.1 |
| StrPur | *Strongylocentrotus purpuratus* | GCF_000002235.5 |
| BraLan | *Branchiostoma lanceolatum* | GCF_035083965.1 |
| CioInt | *Ciona intestinalis* | GCF_018327825.1 |
| HomSap | *Homo sapiens* | GCF_000001405.40 |

Table S10: **Publicly available NCBI short-read archive data used for comparative analyses.**

| SRA | BioProject | Species |
|---|---|---|
| SRR8775109 | PRJNA528777 | *Amphibalanus improvisus* |
| SRR8775110 | PRJNA528777 | *Amphibalanus improvisus* |
| SRR8775111 | PRJNA528777 | *Amphibalanus improvisus* |
| SRR8775112 | PRJNA528777 | *Amphibalanus improvisus* |
| SRR8775113 | PRJNA528777 | *Amphibalanus improvisus* |
| SRR8775114 | PRJNA528777 | *Amphibalanus improvisus* |
| SRR8775115 | PRJNA528777 | *Amphibalanus improvisus* |
| SRR8775116 | PRJNA528777 | *Amphibalanus improvisus* |
| SRR8775117 | PRJNA528777 | *Amphibalanus improvisus* |
| SRR8775118 | PRJNA528777 | *Amphibalanus improvisus* |
| SRR8775119 | PRJNA528777 | *Amphibalanus improvisus* |
| SRR8775120 | PRJNA528777 | *Amphibalanus improvisus* |
| SRR8775121 | PRJNA528777 | *Amphibalanus improvisus* |
| SRR8775122 | PRJNA528777 | *Amphibalanus improvisus* |
| SRR8775123 | PRJNA528777 | *Amphibalanus improvisus* |
| SRR8775124 | PRJNA528777 | *Amphibalanus improvisus* |
| SRR8775125 | PRJNA528777 | *Amphibalanus improvisus* |
| SRR8775126 | PRJNA528777 | *Amphibalanus improvisus* |
| SRR8775127 | PRJNA528777 | *Amphibalanus improvisus* |
| SRR8775128 | PRJNA528777 | *Amphibalanus improvisus* |
| SRR8775129 | PRJNA528777 | *Amphibalanus improvisus* |
| SRR8775130 | PRJNA528777 | *Amphibalanus improvisus* |
| SRR8775131 | PRJNA528777 | *Amphibalanus improvisus* |
| SRR8755479 | PRJNA528169 | *Amphibalanus improvisus* |
| SRR18959849 | PRJNA816681 | *Capitulum mitella* |
| SRR14354747 | PRJNA725059 | *Capitulum mitella* |
| SRR20067441 | PRJNA856037 | *Capitulum mitella* |
| SRR20067442 | PRJNA856037 | *Capitulum mitella* |
| SRR20067443 | PRJNA856037 | *Capitulum mitella* |
| SRR20067446 | PRJNA856037 | *Capitulum mitella* |
| SRR20067447 | PRJNA856037 | *Capitulum mitella* |
| SRR20067448 | PRJNA856037 | *Capitulum mitella* |
| SRR20067449 | PRJNA856037 | *Capitulum mitella* |
| SRR20067444 | PRJNA856037 | *Capitulum mitella* |
| SRR20067445 | PRJNA856037 | *Capitulum mitella* |
| SRR13527601 | PRJNA681321 | *Capitulum mitella* |
| SRR13527602 | PRJNA681321 | *Capitulum mitella* |
| SRR13527603 | PRJNA681321 | *Capitulum mitella* |
| SRR13527604 | PRJNA681321 | *Capitulum mitella* |

### Supplemental Figures

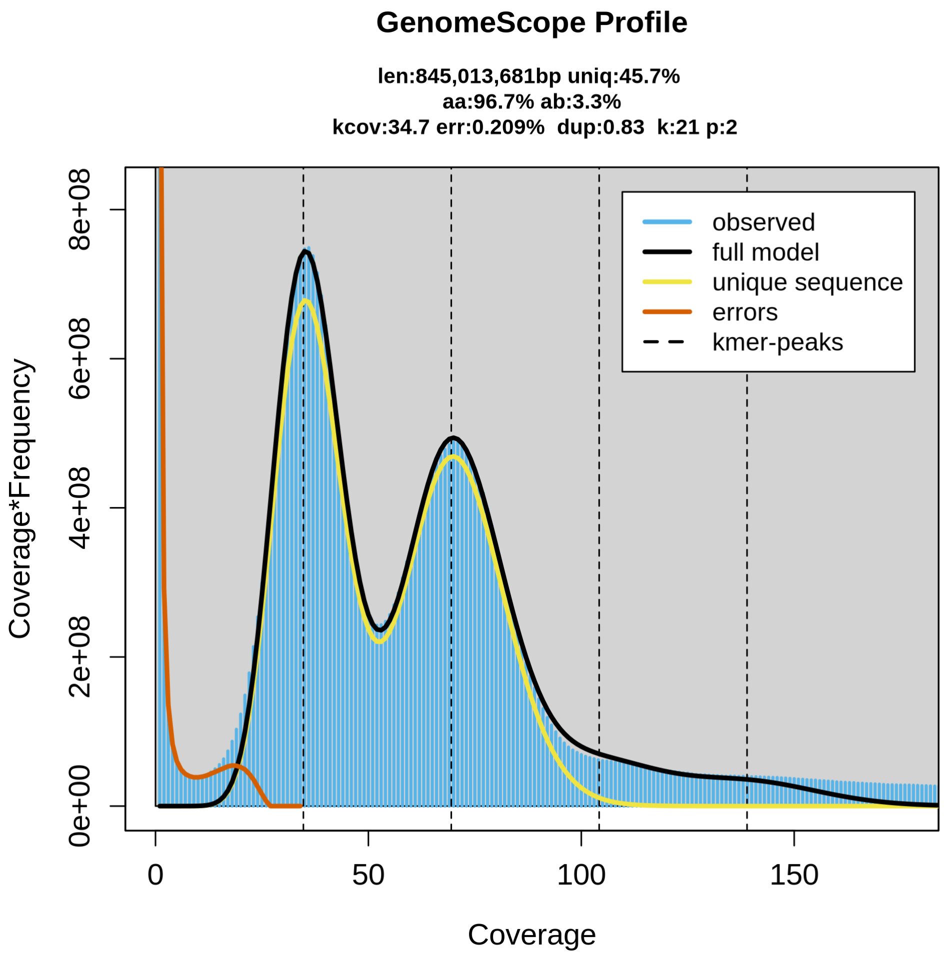


Figure S1: **GenomeScope profile of the raw PacBio HiFi data.** Distribution of 21-mers shows an estimated genome size of 845.01 Mbp and a heterozygosity of 3.3% in the *B. glandula* reference assembly.

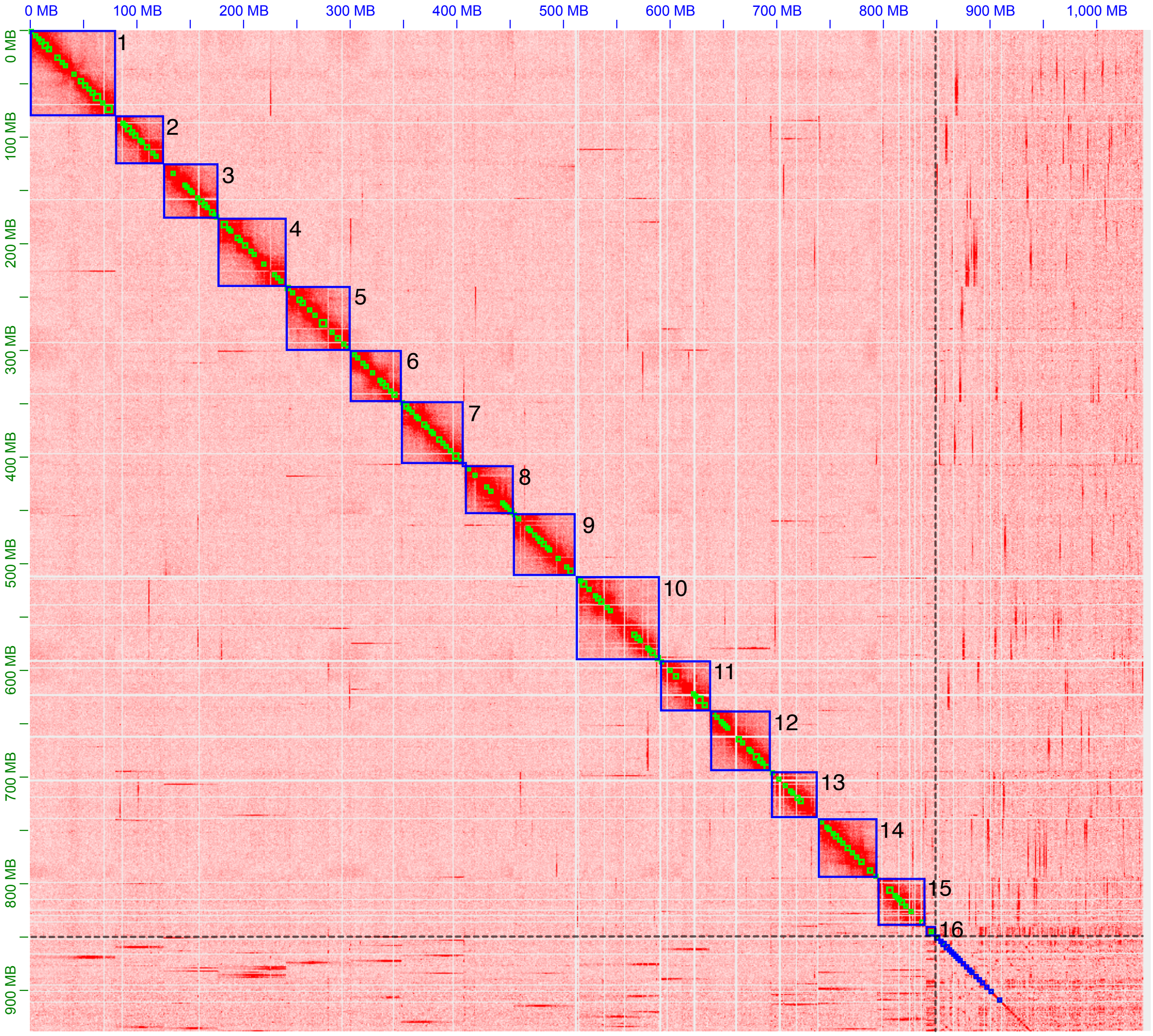


Figure S2: **Hi-C contact map of the *Balanus glandula* assembly.** The contact map shows the assembly following manual curation in Juicebox. The numbers (1–16) denote the 16 chromosome-level scaffolds. The dashed lines show the boundary between these 16 chromosome-level scaffolds and the unplaced contigs. Note that this numbering does not reflect the final ID of the assembled sequences. IDs were re-assigned following additional curation and sorting of the sequences.

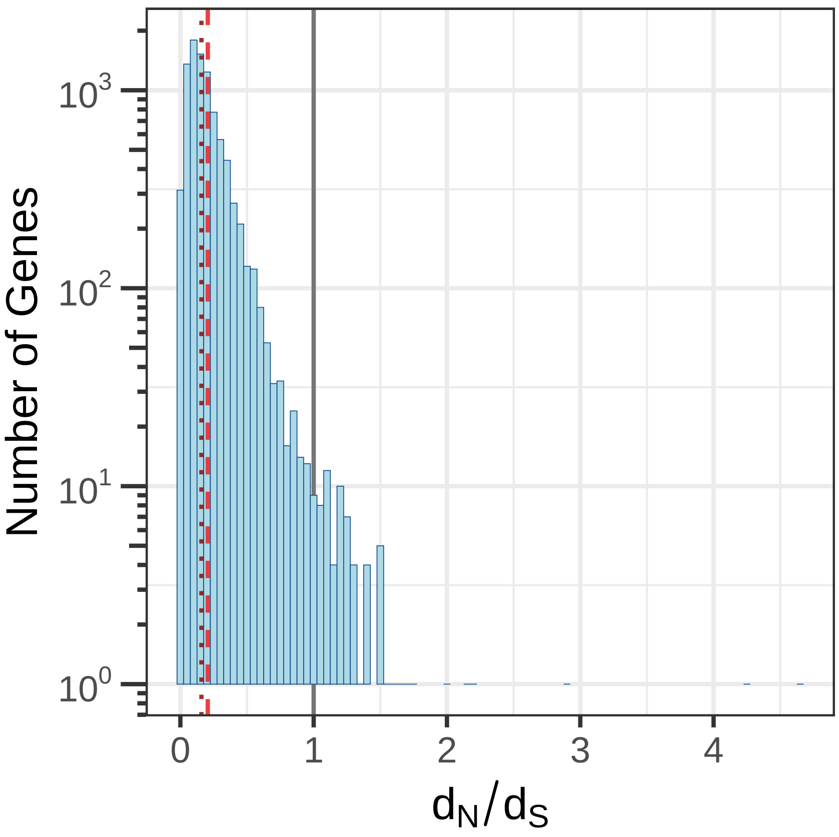


Figure S3: **Distribution of the pairwise rate of non-synonymous to synonymous substitutions ($d_N/d_S$).** The distribution is calculated across 9,082 single-copy orthologs identified between *Balanus glandula* and *Balanus crenatus*. The solid gray line marks the boundary between genes under positive selection ($d_N/d_S > 1$) and genes under purifying selection ($d_N/d_S < 1$). The dashed and dotted lines show the mean (0.206) and median (0.158) $d_N/d_S$, both showing that on average *B. glandula* coding sequences are evolving under a regime of purifying selection.

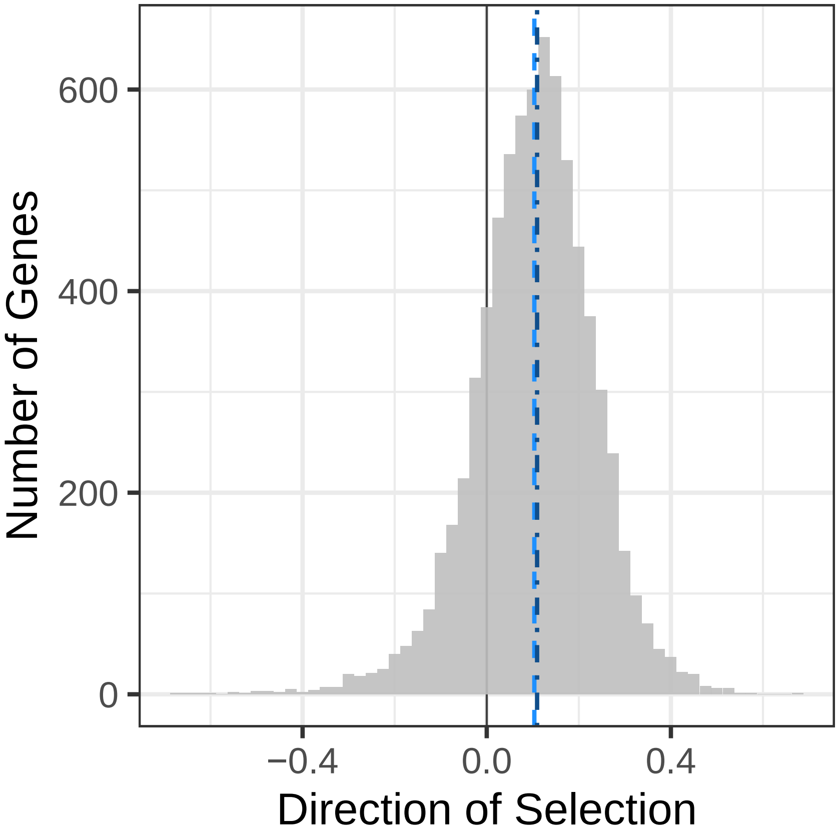


Figure S4: **Distribution of per-gene direction of selection values.** The distribution shows Direction of Selection (DoS) calculated across 7,056 single-copy orthologs identified between *Balanus glandula* and *Balanus crenatus*. The solid black line at zero marks the boundary between genes evolving under a regime of positive selection (DoS $>$ 0) and genes exhibiting an excess of slightly deleterious non-synonymous polymorphism (DoS $<$ 0). The light blue dashed line shows the mean DoS (0.103), while the dark blue dotted line shows the median DoS (0.109). This distribution shows that, on average, coding sequences in *B. glandula* appear to be evolving under positive selection.

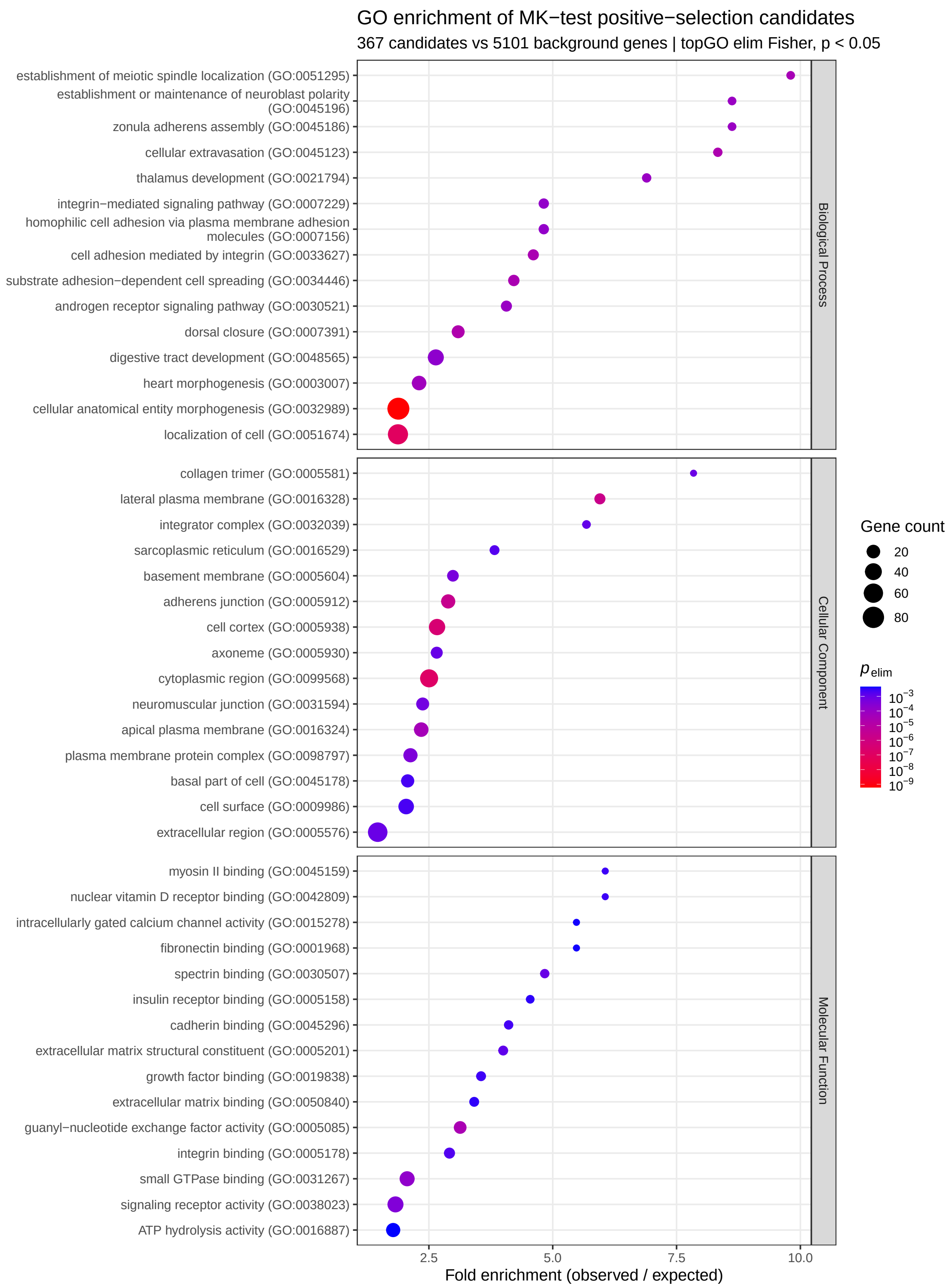


Figure S5: **Significant GO term enrichment among the candidates under positive selection in the McDonald-Kreitman test.** Enrichment for the 367 candidates under selection with annotated GO terms. $p_{\mathrm{elim}}$ denotes the $p$-value calculated from TOPGO's `elim` algorithm. Top, middle, and bottom panels show the top 15 most enriched terms across the three GO categories: biological processes, cellular component, and molecular function.

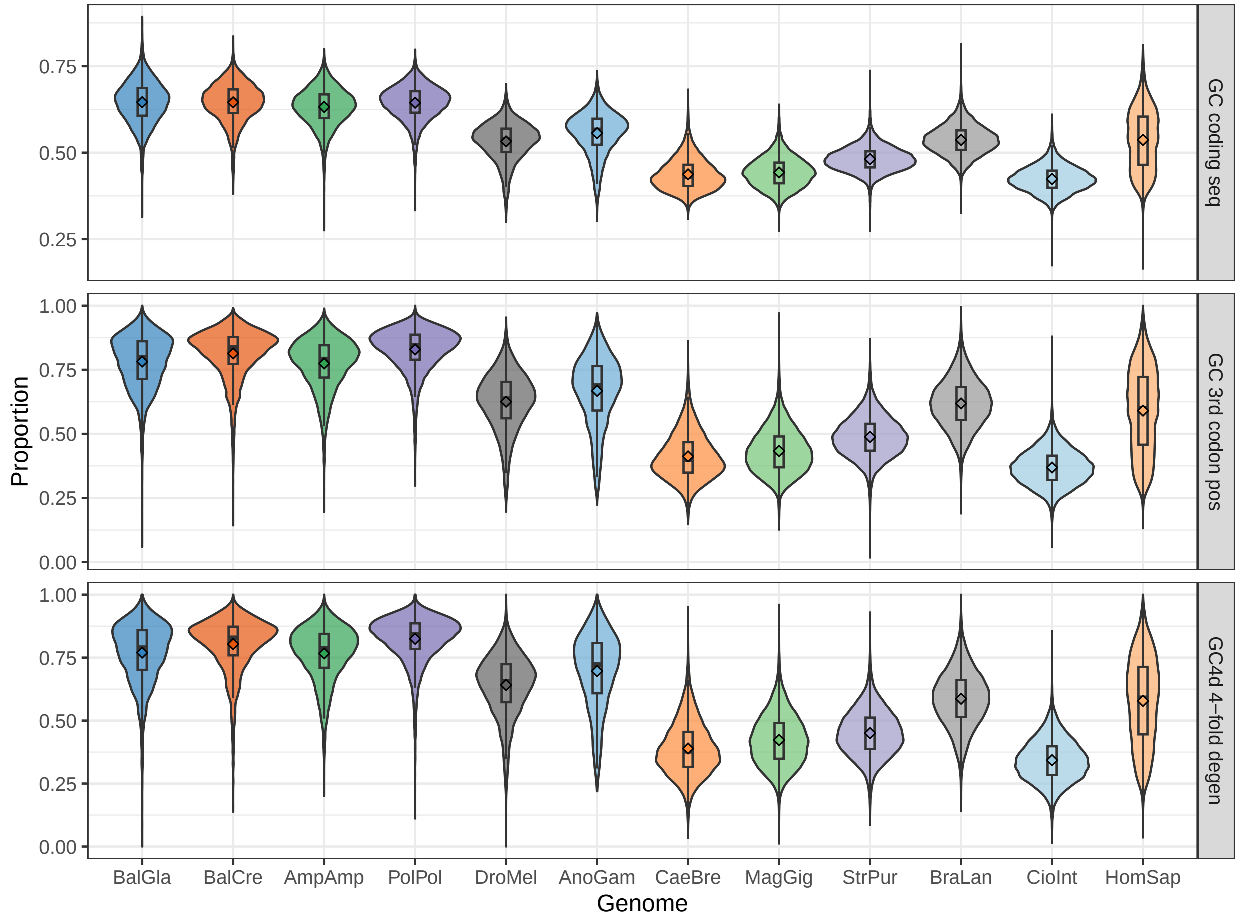


Figure S6: **Distribution of coding GC proportions across metazoan genomes.** Top, middle, and bottom panels show the GC proportions coding sequence-wide, at 3rd codon positions (GC3s), and at 4-fold degerate sites (GC4d), respectively. Distribution are shown across 12 metazoan genomes, including four barnacles: *B. glandula* (`BalGla`), *B. crenatus* (`BalCre`), *A. amphitrite* (`AmpAmp`), and *P. pollicipes* (`PolPol`). See Table S6 for additional description of the genomes.

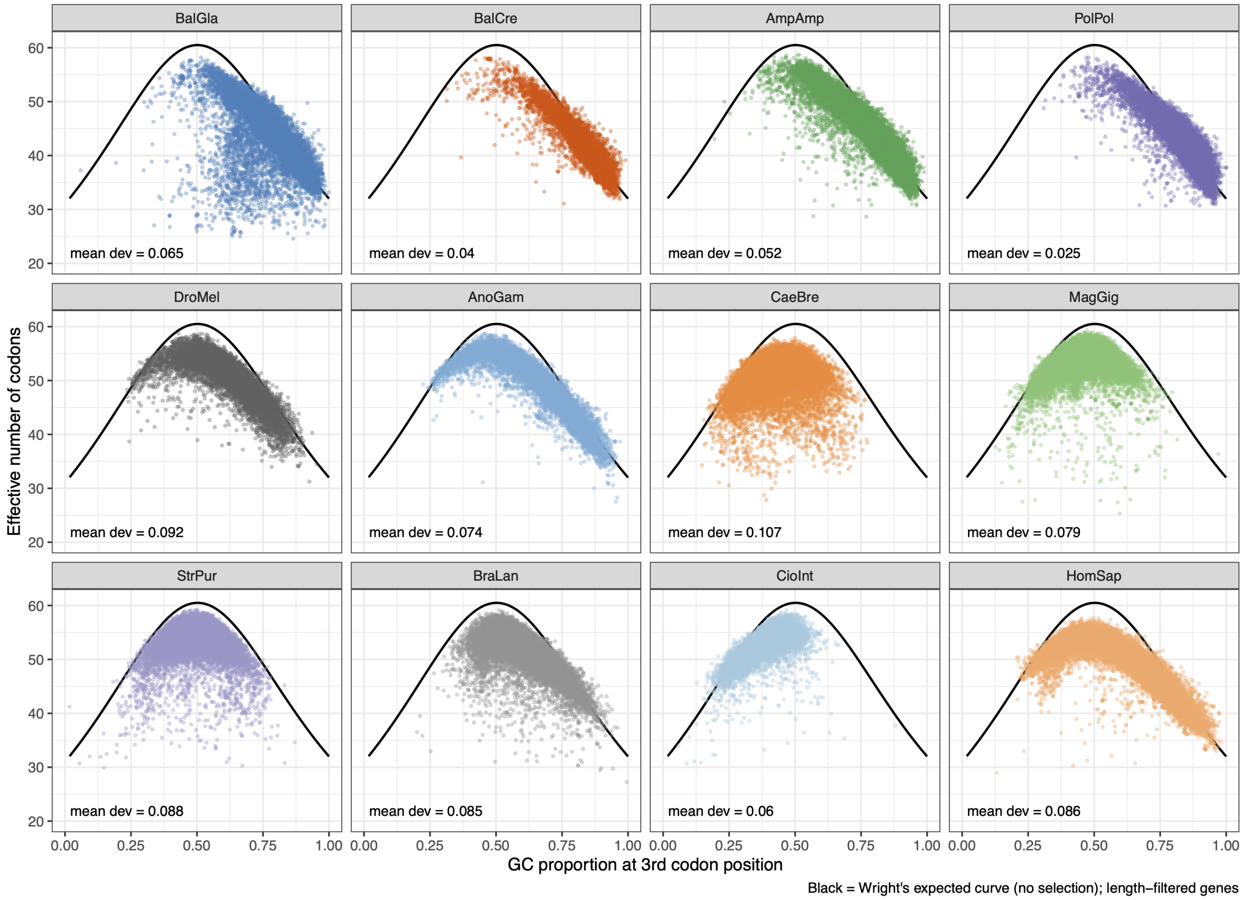


Figure S7: **Relationship between effective codons and GC content.** Nc-plots showing the relationship between GC proportion at 3rd codon positions (GC3s; X-axis) and the effective number of codons (ENC; Y-axis) for 12 metazoan genomes. At each panel, each point represents a single gene. The black curve shows the expected relationship between GC3s and ENC, according to Wright's curve: $ENC_{exp} = 2 + s + 29/[s^2 + (1-s)^2]$, where $s$ denotes GC3s. The "mean dev" annotation shows the mean deviation in ENC from the theoretical expectation. Higher values show a larger proportion of genes showing a smaller than expected number of effective codons, a putative signature of translational selection. See Table S6 for additional description of the genomes.

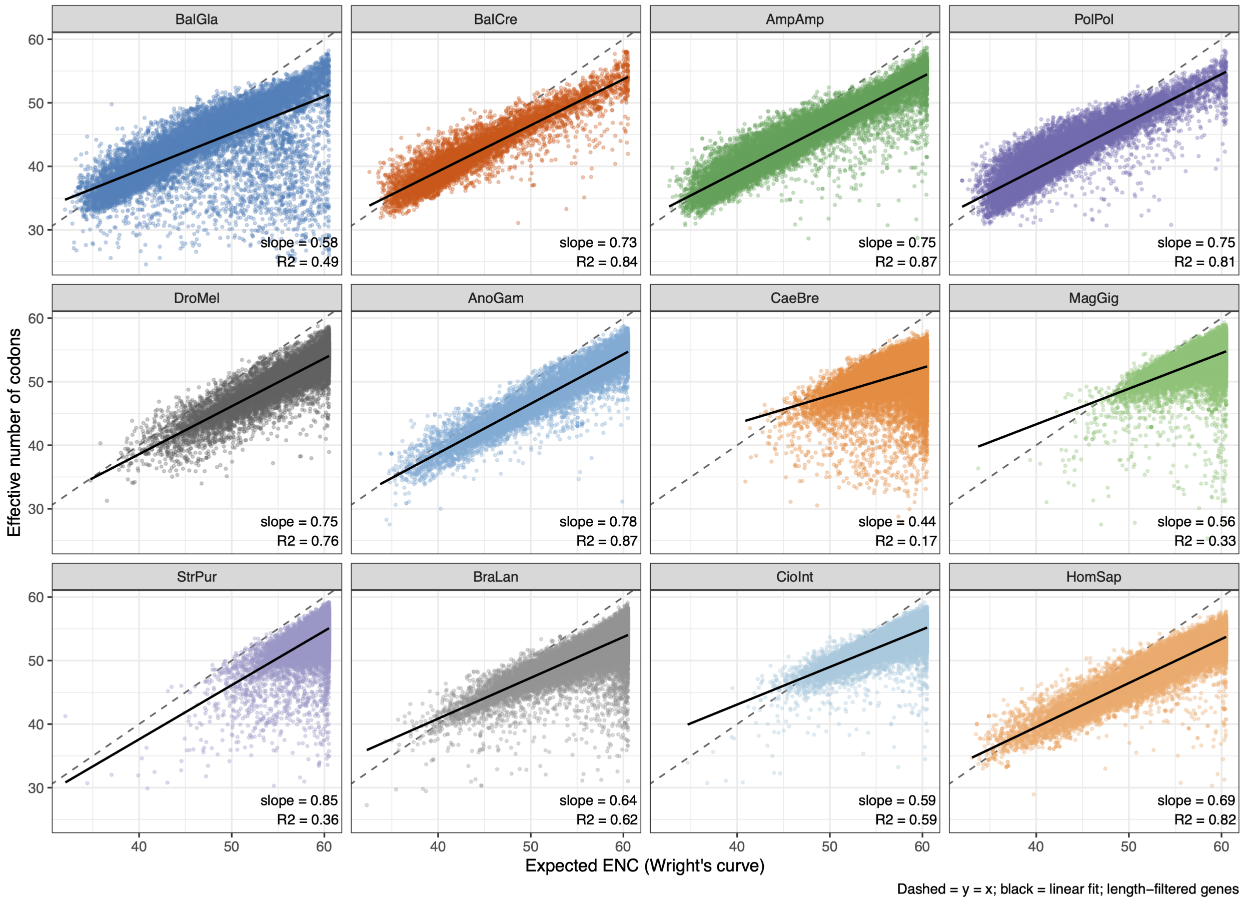


Figure S8: **Relationship between the expected and observed number of effective codons.** Plots show the observed effective number of codons (ENC; Y-axis) as a function of the expected ENC (X-axis), as defined by Wright's curve, for 12 metazoan genomes. At each panel, points represents single genes. Dashed line shows the one-to-one relationship, while the solid black line shows the linear regression. Annotations in the plot show the regression's slope and coefficient of determination ($R^2$). Small ($\ll 1$) slopes and low $R^2$ show a reduction in the observed number of effective codons when compared to the theoretical expectation, a potential signature of translational selection. See Table S6 for additional description of the genomes.

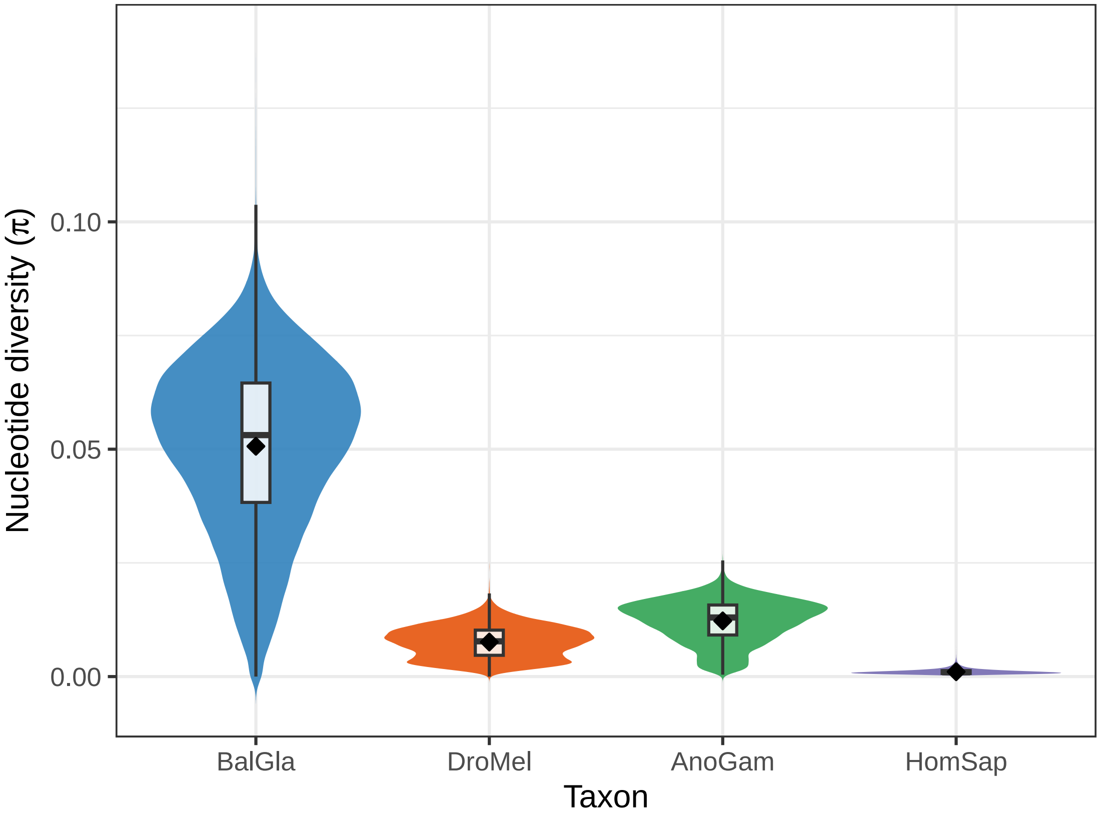


Figure S9: **Comparing nucleotide diversity ($\pi$) between barnacles and biological model systems.** Average $\pi$ in 10 kbp windows between the Oregon Pacific acorn barnacle (*Balanus glandula*; `BalGla`), Sub-Saharan populations of the fruit fly (*Drosophila melanogaster*; `DroMel`), malaria mosquitoes from the Central African Republic (*Anopheles gambiae*; `AnoGam`), and human populations of Yoruba ancestry (*Homo sapiens*; `HomSap`). Consistent with the large $N_e$ expected for *B. glandula*, this barnacle population exhibits an average nucleotide diversity (mean = 0.0506, median = 0.0532, standard deviation = 0.0188) an order of magnitude higher than *Drosophila* (mean = 0.0076, median = 0.0078, standard deviation = 0.0035), $\approx 4\times$ higher than *An. gambiae* (mean = 0.0123, median = 0.0130, standard deviation = 0.0047), and $\approx 48\times$ larger than diverse human cohorts (mean = 0.0011, median = $9.54 \times 10^{-4}$, standard deviation = $5.81 \times 10^{-4}$).

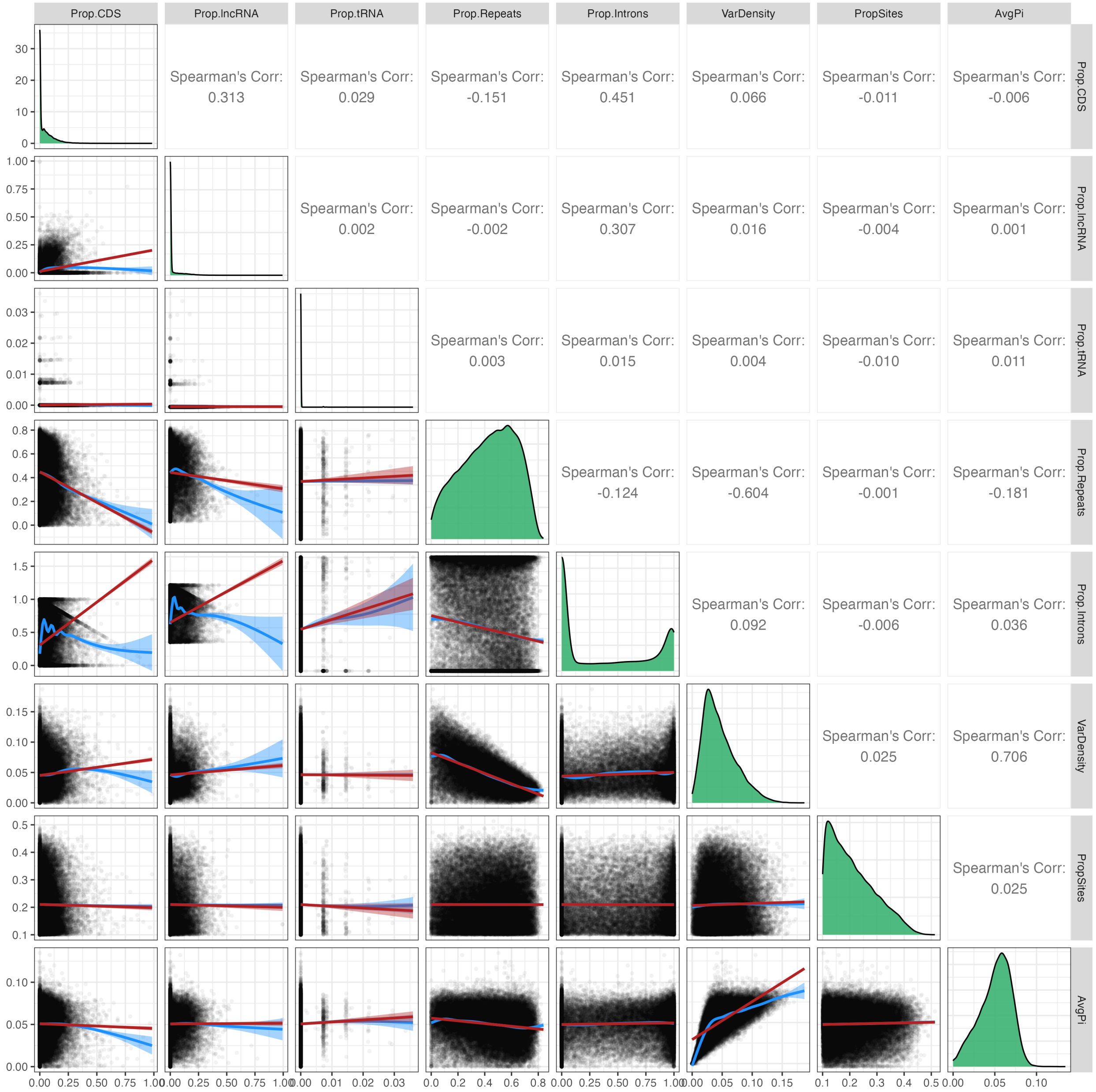


Figure S10: **Pairplot of genomic features and genetic variation in Oregon *Balanus glandula*.** Diagonal shows the density histogram of the proportion of the genomics elements (e.g., CDS, lncRNA, tRNA, repeats, and introns), proportion of variant sites, proportion of callable sites post-filtering, and nucleotide diversity ($\pi$). All values computed across 10 kbp windows. Below the diagonal, plots show the correlation across all pairwise comparisons. Red and blue lines show the linear and Loess regression, respectively. Above the diagonal, panels show the Spearman's rank correlation coefficient ($\rho$) for the given pairwise comparison.

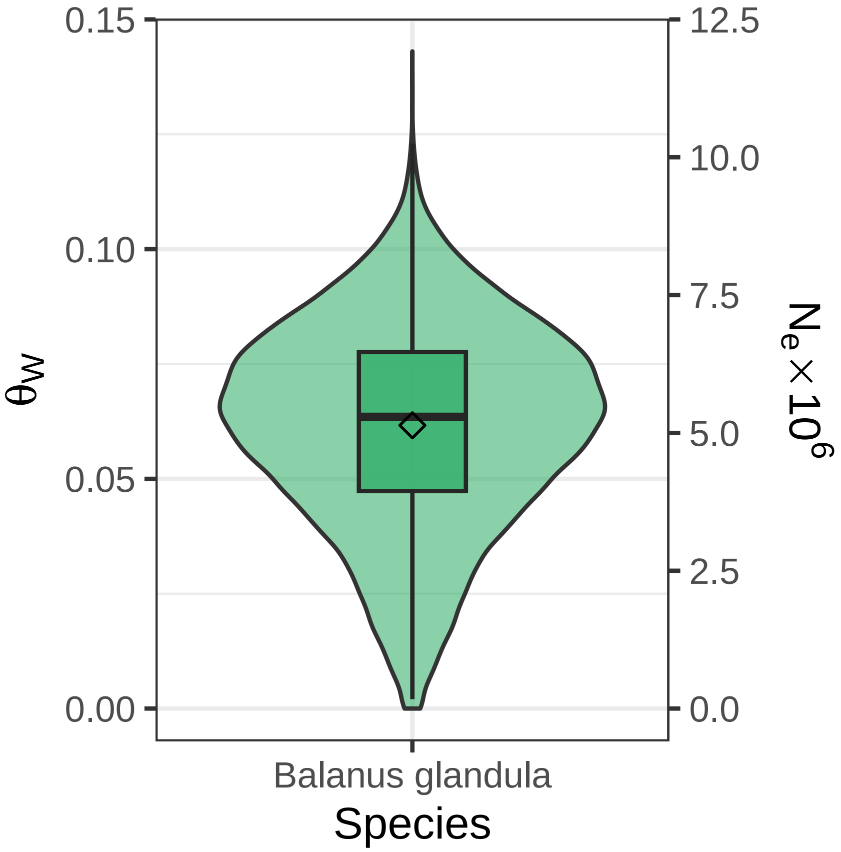


Figure S11: **Effective population ($N_e$) size estimation of *Balanus glandula* based on the Watterson's estimator ($\theta_W$).** Effective size of the population was calculated according to the relationship $\theta_W = 4N_e\mu$, using a mutation rate per-base, per-generation ($\mu$) of $3 \times 10^{-9}$. This estimate yields a long-term $N_e$ of $\approx 5.2 \times 10^6$ for this barnacle population.

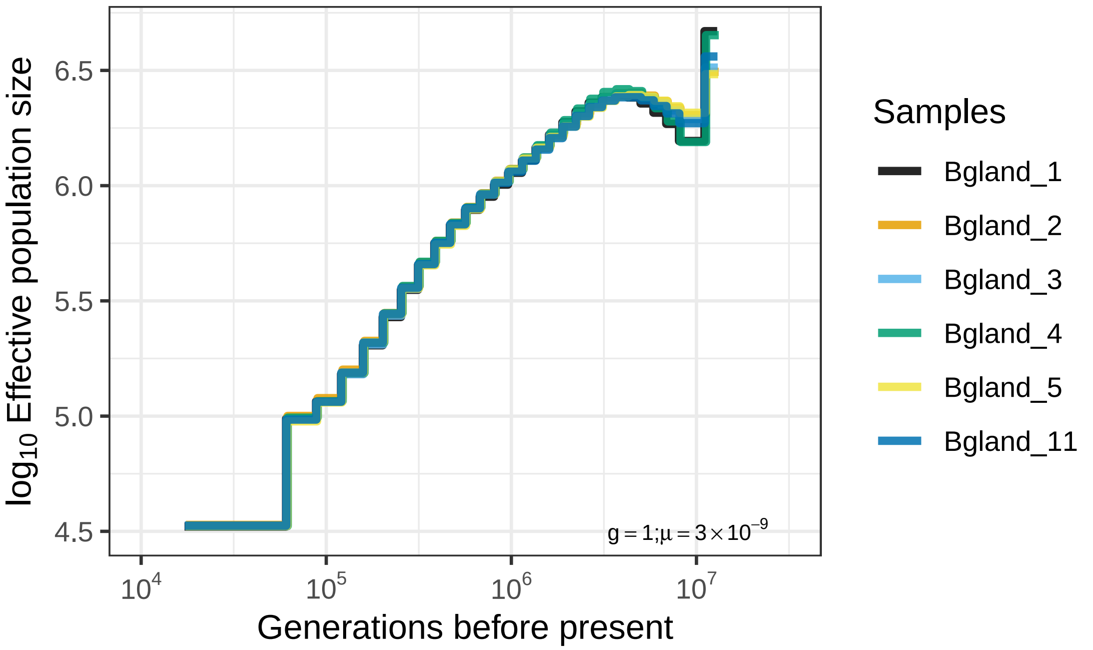


Figure S12: **Coalescent estimation of the effective population ($N_e$) size estimation of *Balanus glandula*.** Estimation of $N_e$ through time was done using the MSMC2 software, using a mutation rate per-base, per-generation ($\mu$) of $3 \times 10^{-9}$, and fixing the generation time ($g$) to 1. Each line represents the $N_e$ trajectory for each sample. Using just six diploid samples, and estimating only between pairs of unphased diploid genotypes within samples, we estimate the historical effective size of the population in the order of $10^6$ individuals; however, we do not obtain enough resolution to resolve $N_e$ at recent ($< 10^5$) time intervals.
